# Interaction between time-of-day and oxytocin efficacy in mice and humans with and without gestational diabetes

**DOI:** 10.1101/2024.11.16.622460

**Authors:** Thu Van-Quynh Duong, Alexandra M. Yaw, Guoli Zhou, Niharika Sina, Aneesh Sai Cherukuri, Duong Nguyen, Kylie Cataldo, Nicollette Ly, Aritro Sen, Lorenzo Sempere, Cara Detrie, Robert Seiler, I. Nicholas Olomu, Rene Cortese, Robert Long, Hanne M. Hoffmann

## Abstract

Management of labor in women with diabetes is challenging due to the high risk of peri- and postpartum complications. To avoid cesarean section and assist with labor progression, Pitocin, a synthetic oxytocin, is frequently used to induce and augment labor. However, the efficacy of Pitocin is often compromised in diabetic pregnancies, leading to increased cesarian delivery. As diabetes deregulates the body’s circadian timekeeping system and the time-of-day of the first Pitocin administration contributes to labor duration, our objective was to determine how the time of day and the circadian clock gene, *Bmal1*, gates oxytocin efficacy. Our studies in mice show that, compared to the rest phase of the day (lights on), the uterotonic efficacy of oxytocin is significantly increased during the active phase (lights off). Using *in vitro* studies, a myometrium-specific *Bmal1* conditional knockout mouse model, and a mouse model of food-induced gestational diabetes, we find that *Bmal1* is crucial for maintaining oxytocin receptor expression and response in the myometrium in mice. These findings also translate to humans, where oxytocin-induced human myometrial cell contraction is time-of-day dependent, and retrospective clinical data suggest that administration of Pitocin in the morning should be considered for pregnant women with gestational diabetes.

## INTRODUCTION

Diabetes during pregnancy represents a critical obstetric concern, contributing significantly to fetal, neonatal, and maternal morbidity and mortality. The incidence of pregestational and gestational diabetes mellitus (GDM) have markedly increased in the United States. GDM rose from 4.6% in 2006 to 8.2% in 2016—a 78% increase within a decade.^1,2^ GDM is associated with an elevated risk of adverse maternal and neonatal outcomes compared to non-diabetic pregnancies. These complications include an increased incidence of preterm birth, prolonged labor, dysfunctional uterine contractions, and postpartum hemorrhage.^3–6^ Notably, women with GDM requiring medication are nearly three times more likely to undergo cesarean delivery.^4^ At the same time, their newborns face a fourfold increase in intensive care unit admissions, and this patient population experiences over five times higher rates of stillbirth compared to newborns of non-diabetic mothers.^3–7^

Diabetes-induced alterations in uterine smooth muscle (myometrium) contractility have been implicated in adverse outcomes.^8–10^ Specifically, the efficacy of Pitocin, a synthetic analog of oxytocin commonly administered to induce and augment labor,^11,12^ is reduced in diabetic pregnancies.^4^ This reduced efficacy is a recognized risk factor for postpartum hemorrhage and emergency cesarean delivery. However, the variability in Pitocin’s effectiveness^13^ prompts further investigation into potential underlying biological factors not yet considered in current clinical management protocols.

One potential contributing factor to the variable efficacy of Pitocin for labor enhancement and labor induction is the time-of-day when Pitocin administration is initiated. We recently identified that Pitocin efficacy was dependent on time-of-day, gestational age, BMI, and parity in non-diabetic pregnant women.^14^ Such time-of-day effects of drugs are well-documented in the literature.^15–18^ One example is Aspirin, a widely used painkiller, and blood thinner with cardiovascular effects. The antihypertensive effect of Aspirin is significantly enhanced when it is taken in the evening compared to the morning.^19,20^ Failure to account for these circadian variations in drug response may result in inconsistent clinical outcomes, requiring higher doses to achieve therapeutic effects, as it has been described for aspirin to prevent preeclampsia.^21–23^

One mechanism by which the time-of-day effects of drugs are generated is through the circadian expression of the drug target. The cell’s endogenous molecular clock, a cell-autonomous transcriptional-translational feedback loop, drives circadian gene expression, which modulates the expression of approximately 20% of cell-specific transcripts,^24–26^ influencing various cellular functions throughout the day.^27–31^ At the core of the molecular clock are a small set of transcriptional regulators, including *Bmal1*, *Clock*, *Cry1*/*2*, *Per1/2/3* and *Nr1d1/2*. Of these transcriptional regulators, loss of *Bmal1* disrupts molecular clock function, leading to a loss in circadian rhythm.^24,25,32^ Dysregulation of the molecular clock has been associated with numerous pregnancy complications in women, such as GDM^33^ and preeclampsia, with and without preterm birth.^34–36^ Notwithstanding, the role of the molecular clock within specific reproductive tissues outside of the ovary and the brain’s suprachiasmatic nucleus remains largely unknown.^37,38^ Several mouse studies indicate the essential role of a functional molecular clock in the uterus during pregnancy.^39–41^ A mouse with a full body dominant negative CLOCK mutation, the DNA binding partner of BMAL1, caused a high rate of fetal resorption and labor dystocia.^42^ In contrast, conditional knockout of *Bmal1* in the smooth muscle (including the myometrium, referred to as *Bmal1* cKO) increased mistimed labor without impacting the transcription level of connexin-3, a contractile-associate protein, oxytocin receptor (*Oxtr*), or progesterone, a hormone reducing uterine contractions, at the studied time points.^43^ Together, these studies indicate that the loss or dysregulation of the molecular clock during pregnancy is associated with poor pregnancy outcomes and potential uterine malfunction.

In this study, we utilized the previously validated *Bmal1* cKO mouse model,^43^ alongside a mouse model of food-induced gestational diabetes (FID) to explore the impact of circadian timing and GDM on oxytocin efficacy. Additionally, in vitro, human myometrial cell studies and retrospective health record data analysis from two hospitals in Michigan were employed to corroborate our findings in humans.

## RESULTS

### Oxytocin uterotonic efficacy depends on the time of day in the gravid mouse

To determine how time-of-day impacts the uterotonic efficacy of oxytocin, we placed uterine explants from late gravid mice (gestation day 18, GD18) in a myograph and recorded their contractile response to oxytocin during the mice’s rest phase (Zeitgeber time 5, ZT5, five hours after lights on ± 3 hours), and active phase (ZT17, five hours after lights off ± 3 hours). Oxytocin increased contraction frequency at both studied time points (Figure 1A). A Two-way ANOVA comparing time-of-day to drug (vehicle and oxytocin) showed a significant effect of time-of-day [F(1, 35)=15.28, p=0.004] and drug [F(1,62)=21.48, p<0.001], but no significant interaction between the two (p=0.68). The effect of oxytocin was specific, as evidenced by the capacity of the oxytocin receptor (OXTR) antagonist, atosiban, to antagonize its effect at both ZT5 and ZT17 (Figure 1A, B). Interestingly, the sole application of atosiban significantly reduced uterine contraction frequency at ZT17 (p=0.0006) but not at ZT5 (p=0.92, Figure 1A). Two-way ANOVA analysis comparing time-of-day to drug (vehicle and atosiban) showed a significant effect of time-of-day [F(1, 42)=4.41, p=0.04), drug (F(1,65)=14.87, p<0.0003), and a significant interaction between the two [F(1,42)=6.77, p=0.01). This suggests that OXTR expression, or the underlying mechanism of uterine contractility, might vary with the time-of-day.

**Figure 1.**
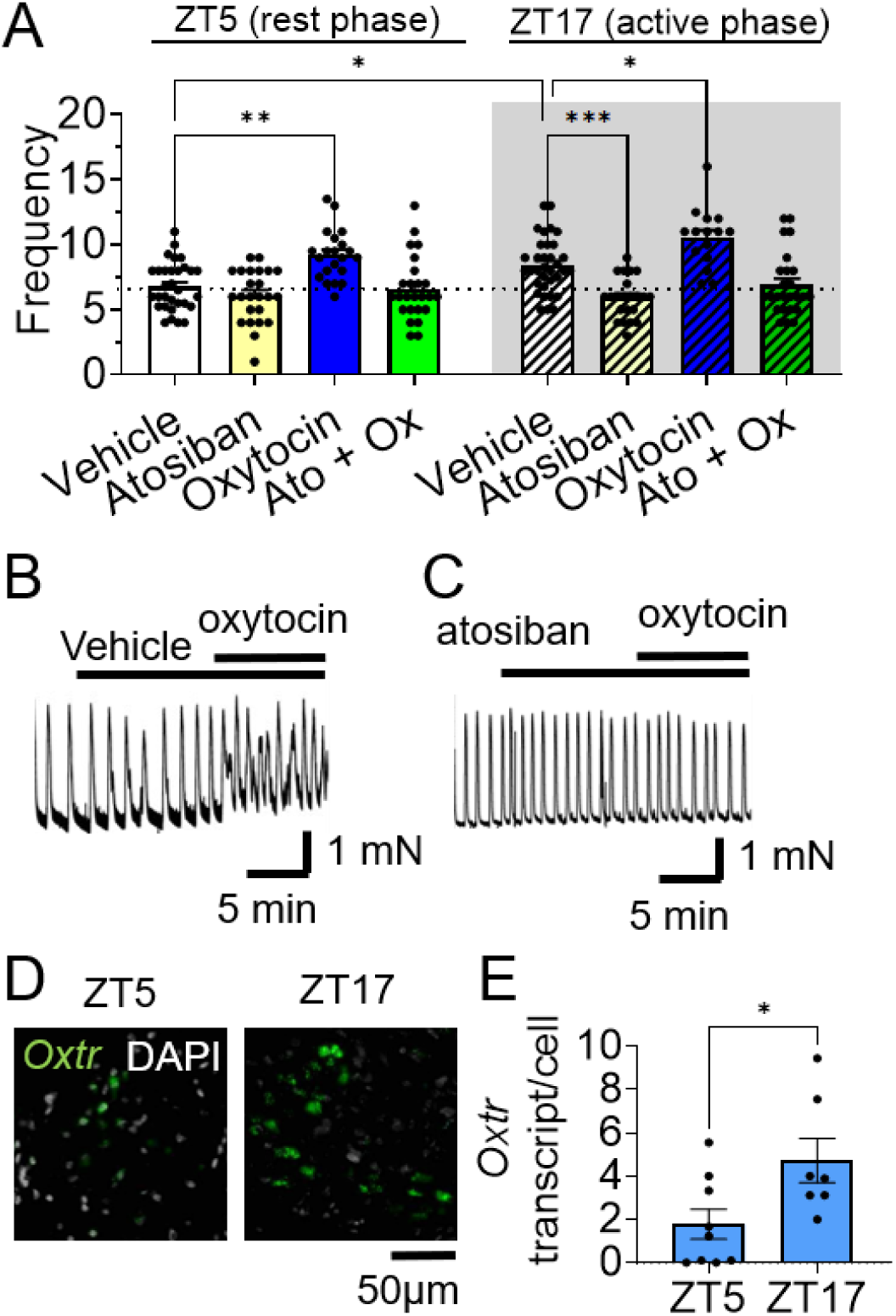
Time-of-day specific effect of OXTR-ligands effect on uterine contractions. **A**) Uterine samples were mounted in a myograph during the dark phase ZT17 (ZT17± hours) or light phase ZT5 (ZT5±3 hours), and contractions evaluated in response to vehicle (water), oxytocin 1nM or atosiban 500nM in wild-type mice at GD18. Two-way ANOVA, n=7-17, with samples in duplicate. *, p<0.05; **, p<0.01; ***, p<0.001. **B, C**) Example tracing of uterine contractions in response to the indicated treatments at GD18, ZT17. **D**) Representative RNAscope® ISH staining for *Oxtr* in the GD18 uterus at ZT5 and ZT17, and **E**) Quantified data. n = 7-9/group. T-test, *, p<0.05.

To determine if the time of day impacted *Oxtr* expression at GD18, we detected *Oxtr* mRNA by the RNAscope® ISH assay. We identified that *Oxtr* was significantly lower during the rest phase of the day (ZT5) than the active phase (ZT17, Figure 1D, E). In contrast*, Bmal1* transcript levels trended lower at ZT17 than ZT5 (t(14) = 2.06, p = 0.059, Supplemental Figure 1A-C), whereas the number of *Oxtr^+^/Bmal1^+^* cells were lower at ZT17 than ZT5 (t(3) = 3.59, p = 0.04, Supplemental Figure 1D-F).

### *Bmal1* is required for circadian rhythms and appropriate oxytocin receptor expression in the mouse myometrium

To further support that the molecular clock gates the time-of-day effect of OXTR-regulated uterine contractions, we conditionally deleted the *Bmal1-flox/flox* allele from smooth muscle cells using the *Telokin-cre+/-* allele. These mice are referred to as *Bmal1* cKO, and have been previously validated.^43^ To confirm the specific abolishment of circadian rhythms in the myometrium, we generated triple transgenic circadian reporter mice, where we crossed the *Bmal1* cKO with the validated PER2::Luciferase^44^ mice, allowing to record PER2-driven circadian rhythms in organotypic tissue explants. We found that *Bmal1* cKO did not impact circadian rhythms in other reproductive tissues than the myometrium (Figure 2A, and Supplemental Figure 2). To determine if BMAL1 targets the *Oxtr* regulatory region, we used the UCSC Genome Browser to identify highly conserved regions of the mouse, rat, human, and monkey *Oxtr* promoter region (Supplemental Figure 3A). Within the conserved regions, we identified four semi-conserved BMAL1 binding sites (E’-boxes, Supplemental Figure 3B). To determine if BMAL1 regulated *Oxtr* expression through these E’-boxes, we performed an *in vitro* reporter gene assay in mouse NIH3T3 cells. We found that Bmal1 overexpression increased Oxtr-luciferase expression in a dose-specific manner (Supplemental Figure 3C). Transient transfections of the Oxtr-luciferase reporter with and without site-directed mutagenesis (labeled μE) to the four E’-boxes identified that mutations at sites μE2 and μE4 abolished Bmal1-induced Oxtr-luciferase expression (Figure 2B). These findings agree with the time-of-day specific binding of BMAL1 to the *Oxtr* regulatory region in male mouse liver (Supplemental Figure 3D). This shows that BMAL1 can directly bind to the *Oxtr* regulatory region to drive *Oxtr* expression. To determine the regulation by BMAL1 of the Oxtr-luciferase reporter *in vivo*, we interrogated *Oxtr* mRNA expression by RNAscope® assay in uterine tissue from *Bmal1* cKO animals (Figure 2C, D). We found that *Oxtr* had a significantly reduced expression in *Bmal1* cKO compared to the control at ZT17 (Figure 2C). It is important to note that *Oxtr* is still expressed in the *Bmal1* cKO myometrium at ZT17, but the number of cells with medium, high, and very high *Oxtr* expression levels is significantly reduced (Figure 2C, p=0.03). Note, due to the low *Oxtr* expression at ZT5 in the control, no difference in *Oxtr* transcript between control and *Bmal1* cKO was identified at this time (not shown), a time point that had a comparable number of *Oxtr* expressing cells in the control and *Bmal1* cKO (p=0.2, Supplemental Figure 4). *Bmal1* was also reduced in the *Bmal1* cKO, in agreement with the previous study validating the targeted cell-type specific genetic deletion in this mouse model (Figure 2D).^43^

**Figure 2.**
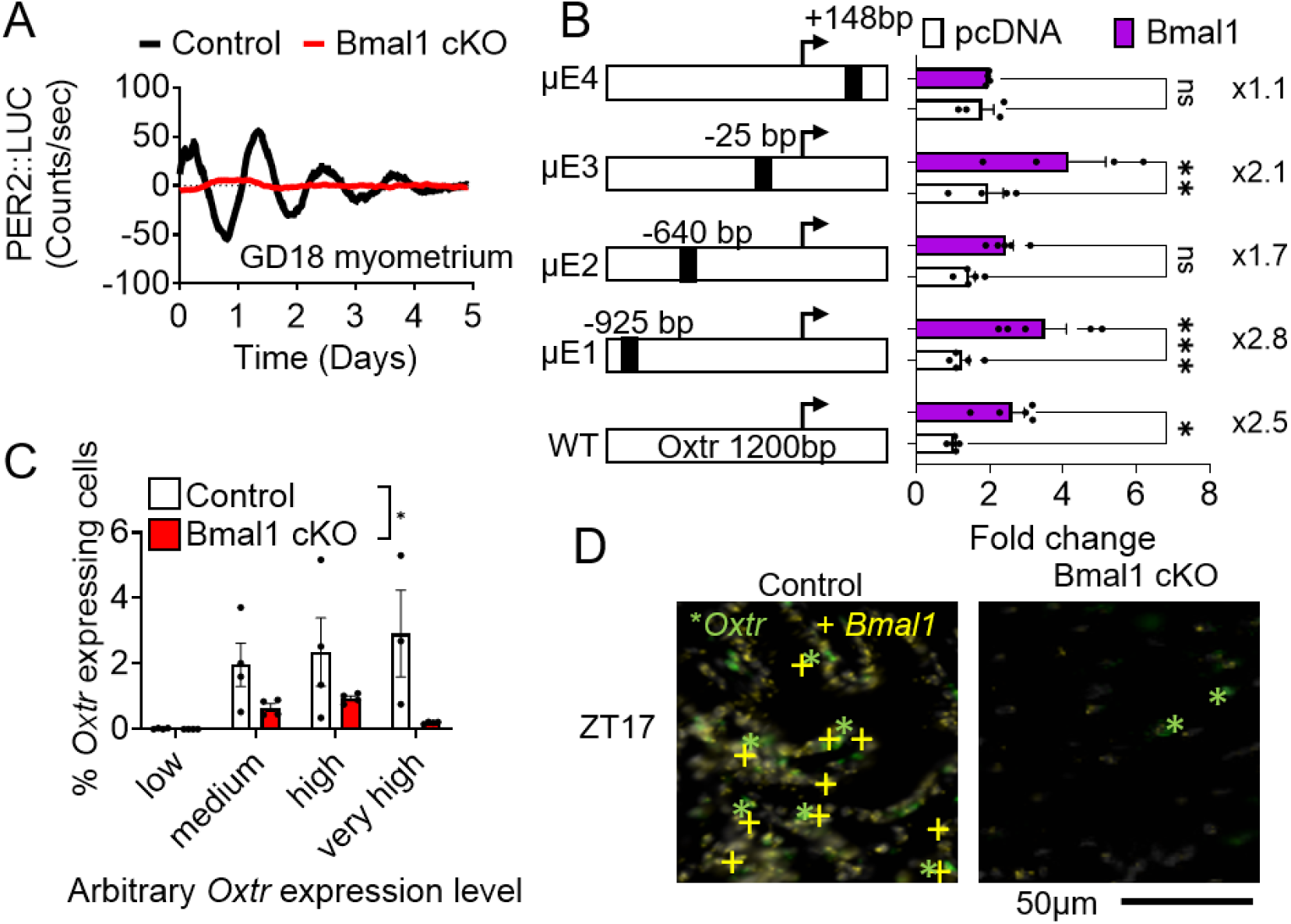
BMAL1 regulates *Oxtr* expression. **A**) Representative PER2::LUC recording of GD18 myometrium from control (Telokin^Cre^:PER2::LUC) and *Bmal1* cKO::PER2:LUC. Data represents n=5/group. **B**) To determine if BMAL1 regulated *Oxtr* expression *in vitro* NIH3T3 cells were transiently transfected with Bmal1 overexpressing plasmid and Oxtr-luciferase plasmids with or without mutated E’-boxes (μE1, μE2, μE3, μE4). Two-way ANOVA, n=4-5 per group in duplicate, ns: non-significant, * p<0.05, ** p<0.05, *** p<0.005. **C**) Histogram of *Oxtr* expression levels and **D**) Representative RNAscope® ISH staining of GD18 uterus at ZT17 in Control and *Bmal1* cKO uterus. Two-way ANOVA, n = 3-4/group. *, p<0.05.

### *Bmal1* cKO uterus is hyposensitive to oxytocin-induced contractions

To test the capacity of oxytocin to promote uterine contractions in *Bmal1* cKO, we first evaluated *ex vivo* spontaneous uterine contraction frequency. Spontaneous uterine contraction frequency was significantly higher in the *Bmal1* cKO than in controls (Figure 3A, F(1,101)=7.26, p=0.008). In addition, in controls, spontaneous uterine contraction frequency was significantly higher during the active phase (ZT17) than in the inactive phase (ZT5, p=0.05), a time-of-day difference lost in the *Bmal1* cKO (Figure 3A, p=0.51). Due to the time-of-day effect on spontaneous uterine contraction frequency, we normalized the oxytocin-induced contraction data to vehicle control within each sample for the time points studied (Oxytocin/Vehicle). As shown in Figure 1A, oxytocin increased uterine contractions independent of the time of day in controls. Interestingly, the capacity of oxytocin to increase uterine contraction frequency was significantly reduced at ZT5 in *Bmal1* cKO as compared to control (Figure 3B, C, p=0.002), with an overall reduced efficacy of oxytocin in the *Bmal1* cKO (p<0.01). Together, these results indicate that loss of *Bmal1* expression in the mouse myometrium causes a reduction in *Oxtr* expression (Figure 2C, D), which is associated with a reduced capacity of oxytocin to increase uterine contraction frequency (Figure 3B).

**Figure 3.**
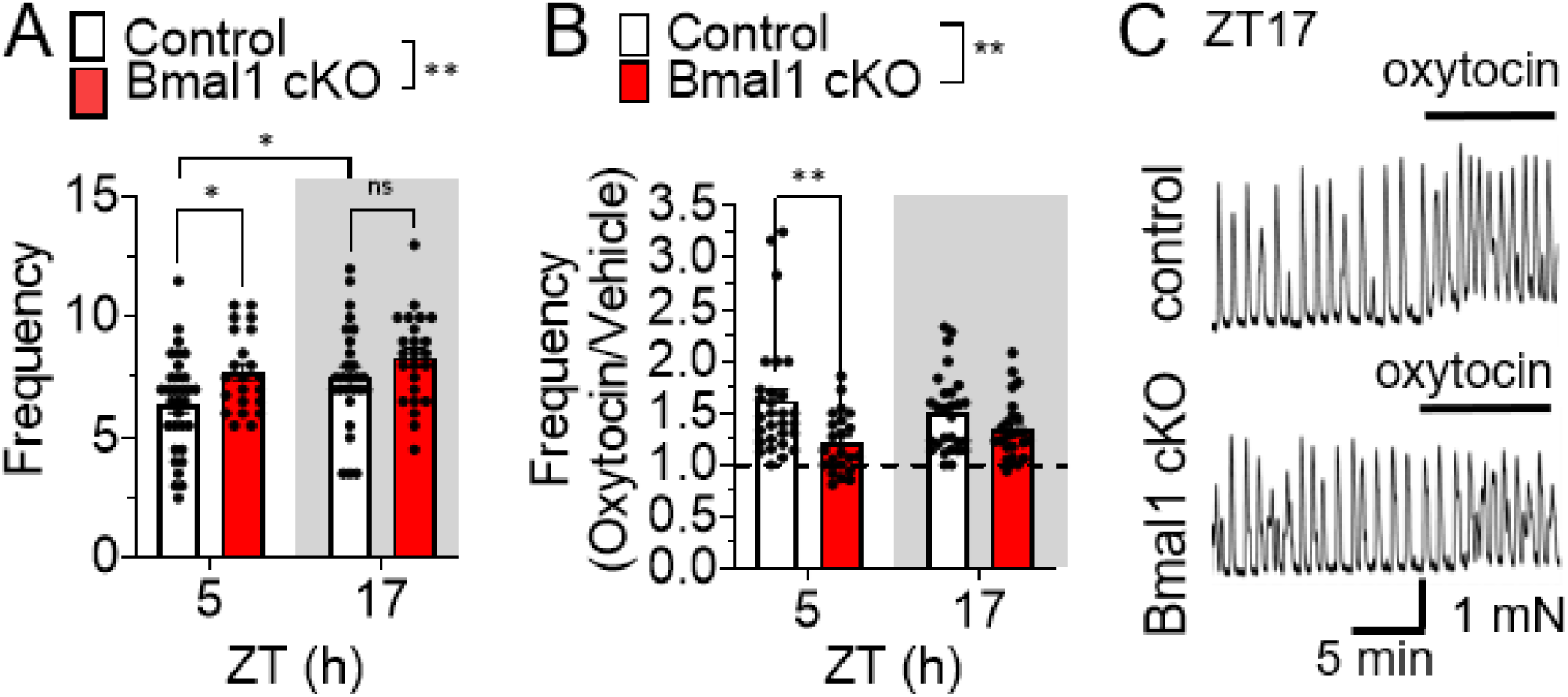
Time-of-day and *Bmal1* cKO impacts oxytocin’s uterotonic efficacy in mice. **A**) GD18 uterine samples were mounted in a myograph, and spontaneous uterine contraction frequency was analyzed at ZT5 and ZT17 in control and *Bmal1* cKO. n = 11-15, in duplicate. Two-way ANOVA, **, p<0.01. **B**) Histogram shows the effect of oxytocin 1nM-promoted uterine contraction frequency normalized to baseline frequency in GD18 uterine samples harvested at ZT5 and ZT17 from control and *Bmal1* cKO. Two-way ANOVA, n = 11-14, in duplicate. **, p<0.01. **C**) Example uterine contraction recordings in response to oxytocin at GD18, ZT17 in control and *Bmal1* cKO.

### Food-induced gestational diabetes (FID) reduced oxytocin-induced contractions and molecular clock transcript expression in the mouse uterus

GDM is known to deregulate the molecular clock and reduce oxytocin efficacy. Based on this, we asked how GDM in the mouse impacted molecular clock expression and oxytocin’s efficacy in promoting contractions in the mouse. To generate females with GDM, we placed virgin females on standard chow or high fat, high sucrose diets for 5 weeks prior to pregnancy. Females were maintained on the assigned diets throughout the study (Figure 4A). All high fat, high sucrose diet mice had their fasting glucose levels tested and responses to a glucose tolerance test (GTT) completed on GD16. Based on the GTT, females were labeled as Food-normal GTT (FN) if they had a normal GTT (GTT<180mg/ml) and as Food-induced gestational diabetes (FID) if they failed the GTT (GTT>180mg/ml, Supplemental Figure 5A, B). Both FN and FID females gained more weight over the study period than females on standard chow (Supplemental Figure 5C). In addition, the GD19 fetuses from mothers in the FN and FID groups had significantly higher blood glucose and insulin levels compared to GD19 fetuses from Ctr mothers on standard chow (Supplemental Figure 5D, E).

**Figure 4.**
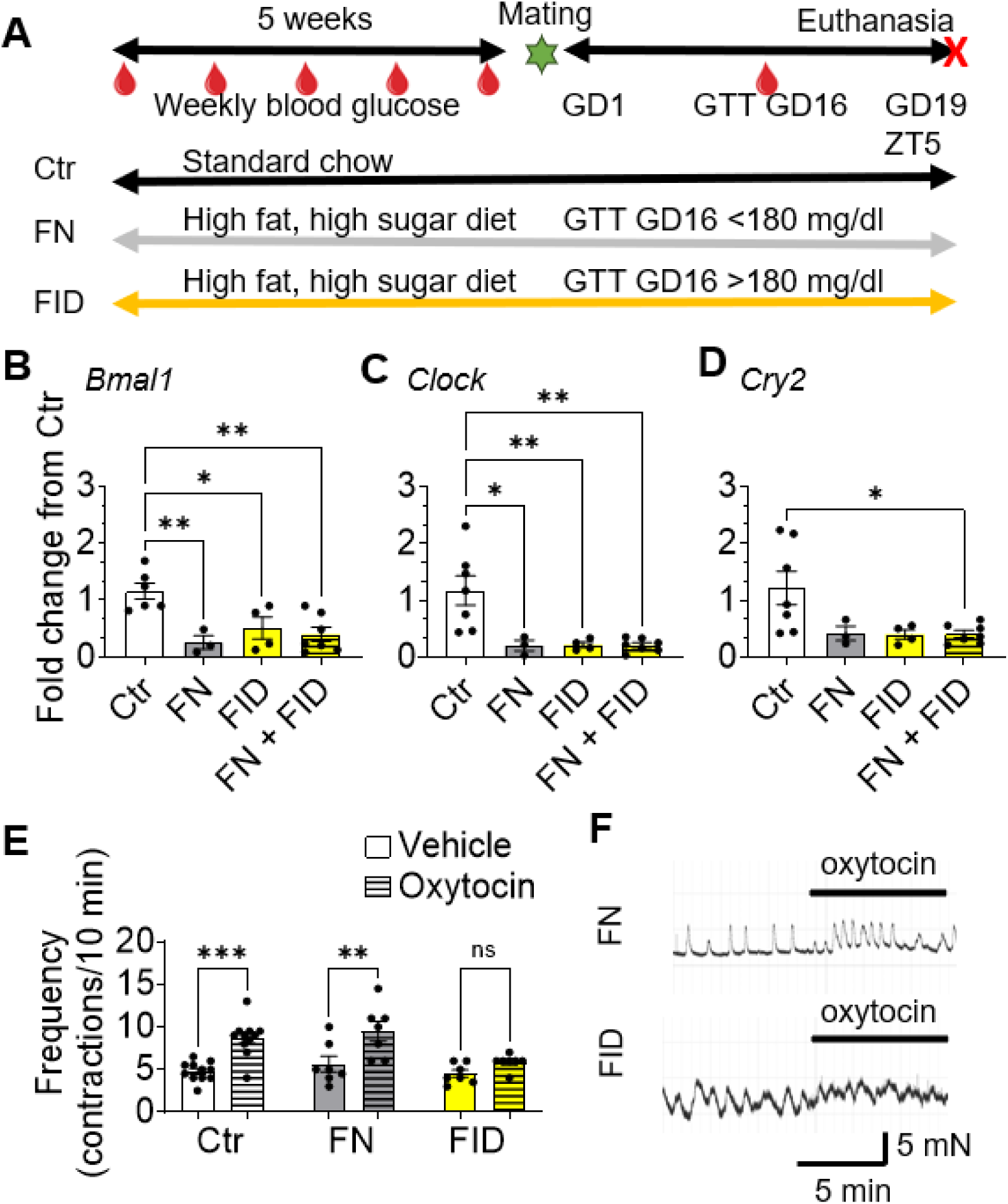
Food-induced gestational diabetes (FID) reduces molecular clock expression and the uterotonic efficacy of oxytocin. **A**) An experimental timeline for generating Control (Ctr), Food-normal GTT (FN), and FID mice. **B-D**) qPCR of the uterine sample at GD18, ZT5. One-way ANOVA, n = 3-7/group, *, p<0.05; **, p<0.01. **E)** Histogram, **F)** example recording of uterine contractions in response to vehicle or oxytocin 1nM at GD18, ZT5. Two-way ANOVA, n=3-5, with samples in duplicate. *, p<0.05; **, p<0.01; ***, p<0.001.

To determine how the high fat, high sucrose diet with (FID) and without (FN) GDM impacted molecular clock gene expression, we collected GD19 ZT5 uterine samples for qPCR. Based on previous work showing that these transcripts are deregulated in the blood and placenta of women with pregestational diabetes, preterm birth, and preeclampsia, we focused on *Bmal1*, *Clock*, and *Cry2*.^33–36^ We found that the uterus of females on standard chow (Ctr) had significantly higher levels of *Bmal1* and *Clock* than FN, FID and FN + FID (FN and FID were merged as they were not significantly different, Figure 4B, C). BMAL1-CLOCK drives the expression of the clock gene *Cry2*, a transcriptional regulator significantly lower in the FN + FID group than Ctr (Figure 4D). To determine how the reduction in molecular clock transcripts in the FN and FID mice was associated with the capacity of oxytocin to promote uterine contractions, we placed GD19 ZT5 uterine explants on the myograph. No spontaneous uterine contraction frequency difference was observed for vehicle across different groups (Ctr, FN, and FID, Figure 4E). In contrast, FN uterine samples responded with increased uterine contraction frequency to oxytocin, while the FID group was associated with a loss of uterotonic efficacy of oxytocin (Figure 4E, F).

### Human myometrial cells have time-of-day-specific responses to oxytocin

Our recent work identified that in pregnant women without diabetes, the time-of-day impacted Pitocin (synthetic oxytocin)-induced labor duration.^14^ To determine if this time-of-day effect of Pitocin might arise at the level of myometrial cells, we tested *in vitro* how time-of-day impacted the uterotonic efficacy of oxytocin. We used a validated protocol to synchronize the molecular clock in cells^45,46^ (Figure 5A) and assessed the capacity of oxytocin to promote term human fibroblast-like myometrial cell (PHM1-41 cell^47^) contractions. Vehicle (water) did not impact PHM1- 41 cell size (used as a measure of contraction); in contrast, oxytocin-induced PHM1-41 cell contractions in a time-of-day dependent manner, with a significant effect on cell contraction (% cell size) at circadian time points 0, 4, 10, and 13 hours (Figure 5B).

**Figure 5.**
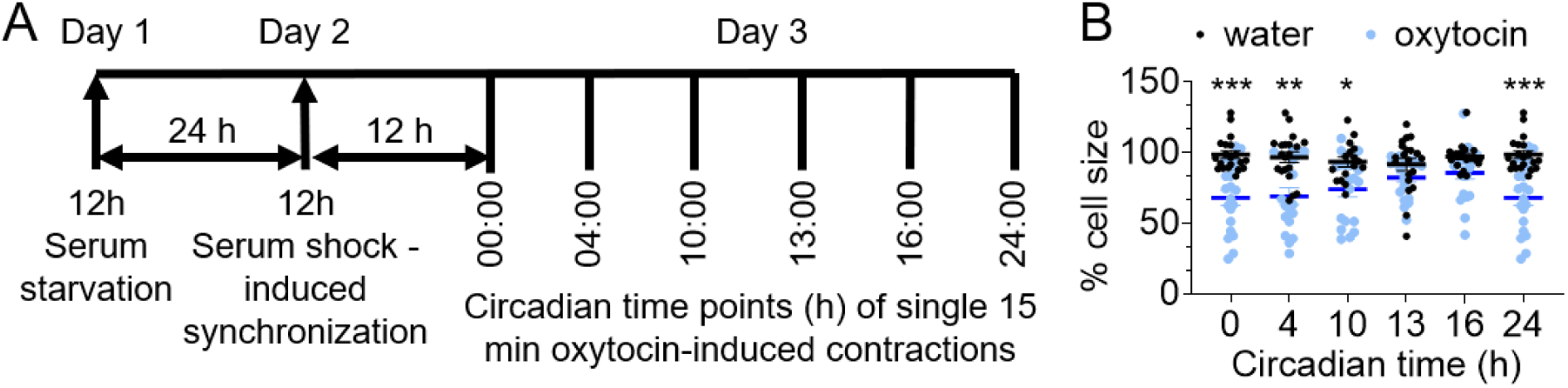
Human myometrial cells exhibited time-dependent contractile response to oxytocin. **A**) Experimental timeline of synchronization and treatment of PHM1-41 cells. After treatment with serum-free media for 24 hours (serum starvation), cells were treated with regular cell culture media to synchronize their circadian rhythms. Beginning at 12 hours after synchronization (time point 0:00 hours), each culture dish was treated with either vehicle (water) or 4.4 nM oxytocin for 15 minutes at the indicated time points. Images were captured at 0 and 15 minutes after treatment and analyzed for cellular surface area to measure cellular contraction. **B**) Changes in PHM1-41 cell size (% from 0:00 hours) to oxytocin at indicated time points. Two-way ANOVA with mixed effects, n=3 replicates per time point with 5-10 cells/replicate, *p<0.05, **p<0.01, ***p<0.001.

### Time-of-day of labor enhancement and labor induction impacts labor duration in a GDM- specific manner

Pregestational and gestational diabetes in women are associated with down and dysregulated molecular clock expression.^33^ To determine if this known effect of diabetes on the body’s circadian clock mechanism impacted the labor duration in a time-of-day-specific manner, we used retrospective data from two independent patient cohorts. As summarized in Tables S1 and S2, the Sparrow cohort (n= 7,804) consisted of pregnant women with 5.2% Hispanic ethnicity, 44.9% maternal age ≥30 years old, 52.2% parity≥1, 66.7% obesity (BMI>30 kg/m^2^), and 5.2% GDM occurrence. The McLaren cohort (n= 1,632) consisted of patients with 5.1% Hispanic ethnicity, 30.7% maternal age≥30 years old, 28.6% parity>1, 41.1% obesity, and 9.5% GDM occurrence. To determine how the time-of-day of initiation of labor impacted labor duration, we defined labor duration as the time-of-day the first labor induction procedure was initiated. Women with and without assisted cervical ripening (Cytotec/Misoprostol and Foley catheter) were included, where 100% of the patients received Pitocin, with 56% of the patients in the Sparrow dataset receiving Cytotec/Misoprostol and 8% Foley catheter, whereas 100% of the women in the McLaren dataset received Cytotec/Misoprostol before Pitocin. The induction time was defined as the initial dose of a cervical ripening agent or Pitocin, whichever was first. Using these definitions, bivariate association analyses demonstrated that without the adjustment of other significant covariates (e.g., maternal age, parity, and BMI), induction time-of-day was significantly associated with labor duration, with the longest labor durations in 0:00-4:00 and 20:00-24:00 hours intervals (average labor duration=19.1 ± 11.5 hours for 0:00-4:00 and 19.3 ± 12.0 hours for 20:00-24:00 hours) and the shortest labor durations in 4:00-8:00 and 8:00-12:00 hours intervals (average labor duration=15.8 ± 9.0 hours for 4:00-8:00 hours and 17.0 ± 9.7 hours for 8:00-12:00 hours, with an overall effect of time of induction p<0.0001) in the Sparrow cohort (Table S1, Figure 6). Compared to non-GDM, GDM had significantly longer labor duration in the Sparrow cohort (average labor duration in GDM=20.6 ± 13.0 hours and in non-GDM=18.2 ± 11.0 hours, p=0.0035, Table S1). Although limited by its smaller sample size, similar association patterns were found for the McLaren cohort. Specifically, overall, the induction time (Table S2, p=0.0001), and BMI (Table S2, p=0.0043) impacted labor duration. Whereas, the relationship of induction time and GDM with labor duration, though not statistically significant, was longest in 20:00-24:00 hours intervals (average labor duration=18.4 ± 8.9 hours) and shortest in the 4:00-8:00 hours and 8:00-12:00 hours intervals (average labor duration=13.4 ± 5.2 to 13.5 ± 7.0 hours; average labor duration in GDM=15.8 ± 7.5 hours and in non-GDM=14.7 ± 8.0 hours, p=0.310, Table S2, Supplemental Figure 8). Multiple linear regression analyses revealed significant main effects (including directions) of induction time in non-GDM patients in the Sparrow cohort after adjusting for parity and BMI. Labor duration significantly decreased by 3.34 ± 0.78 hours in non-GDM patients when induction was conducted during the 0:00-4:00 hour interval compared with 4:00-8:00 hour interval (p<0.0001). Furthermore, labor duration in non-GDM patients was significantly reduced by 1.61 ± 0.66 hours when induction was conducted at 0:00-4:00 interval compared with 8:00-12:00 hour interval (p=0.0147). Overall, GDM led to an average increase of 1.85 ± 0.74 hours in labor duration compared to non-GDM (p=0.0123, Table 1). The interaction between induction time and GDM significantly contributed to the labor duration in the Sparrow cohort (Table 1). To obtain complete multiple comparisons for the interaction term in a multiple linear regression model, we estimated the marginal means of labor duration across different induction intervals with and without GDM in the Sparrow cohort (see specific groups in Tables 2 and 3). The results show that the significantly increased labor duration in GDM patients compared to the non-GDM peers only occurred in the 0:00-4:00 hours induction time interval (estimated marginal mean of labor duration=10.25 ± 1.84 hours, adjusted p<0.0001, Table 2, Figure 6). Meanwhile, multiple comparisons of the estimated marginal means of labor duration across all induction time intervals within GDM or non-GDM revealed that induction time intervals in patients without GDM were significantly different between 0:00-4:00 vs 4:00-8:00 hours (p=0.0001), 4:00-8:00 vs 16:00-20:00 hours (p<0.0001), 4:00-8:00 vs 20:00-24:00 hours (p<0.0001), 8:00-12:00 vs 16:00-20:00 hours (p=0.009) and 8:00-12:00 vs 20:00-24:00 hours (p=0.01). In patients with GDM labor duration was significantly different between 0:00-4:00 vs 4:00-8:00 hours (p=0.0005); 0:00-4:00 vs 8:00-12:00 hours (p=0.022); 0:00-4:00 vs 12:00-16:00 hours (p=0.0005); 0:00-4:00 vs 16:00-20:00 hours (p=0.0005); and 0:00-4:00 vs 20:00-24:00 hours (p=0.0091, Table 3). In the McLaren cohort, GDM’s main and interaction effects of induction time with GDM trended in the same direction as in the Sparrow dataset (Tables S3-S5, Supplemental Figure 8). Still, statistical significance was not reached, probably due to a lower sample size.

**Figure 6.**
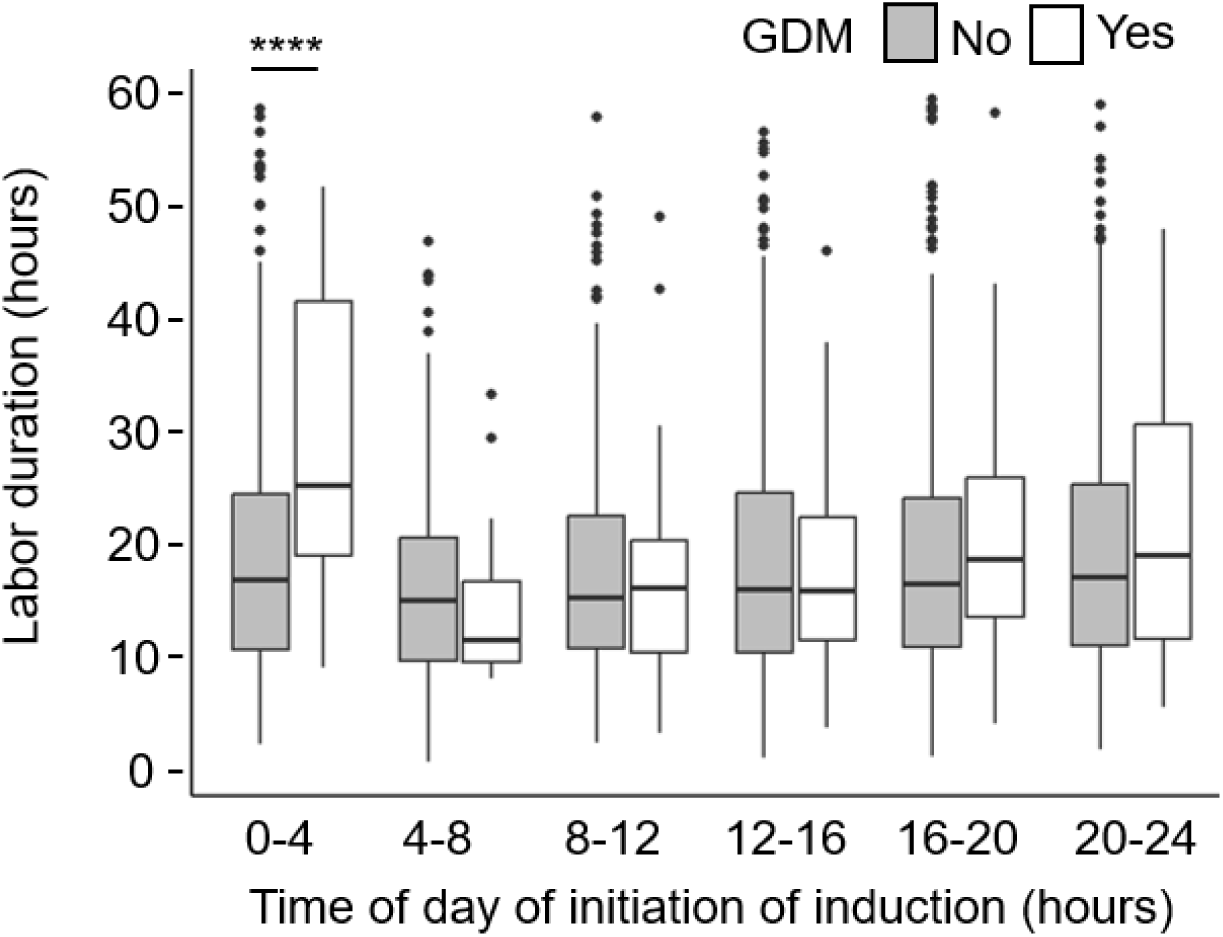
Labor duration as a function of the time-of-day of labor induction in women with and without GDM. The average labor duration for each 4-hour time of induction bin across the 24 hour day is summarized for women with (Yes, n=187) and without (No, n=3426) GDM in the Sparrow cohort. Linear regression with a comparison of labor duration between GDM yes (white boxes) vs. no (gray boxes), ***p<0.001. For complete statistics, see Tables 2 and 3.

**Table 1.** Summary of the association between the outcome variable Labor Duration and the covariates (Sparrow, N=3613).

| | Coefficient ($\beta$) | SE of $\beta$ | t Statistic | p |
| --- | --- | --- | --- | --- |
| **Intercept:** | 19.73 | 0.80 | 24.52 | <0.0001 |
| **Time of Induction:** |  |  |  |  |
| 4-8 vs. 0-4 | -3.34 | 0.78 | -4.30 | <0.0001 |
| 8-12 vs. 0-4 | -1.61 | 0.66 | -2.44 | 0.0147 |
| 12-16 vs. 0-4 | -0.36 | 0.58 | -0.63 | 0.5298 |
| 16-20 vs. 0-4 | 0.18 | 0.58 | 0.31 | 0.7568 |
| 20-24 vs. 0-4 | 0.11 | 0.58 | 0.19 | 0.8463 |
| **GDM:** |  |  |  |  |
| Yes vs. No | 1.85 | 0.74 | 2.50 | 0.0123 |
| **Parity:** |  |  |  |  |
| $\geq 1$ vs. 0 | -9.06 | 0.33 | -27.29 | <0.0001 |
| **BMI:** |  |  |  |  |
| Overweight vs. Normal | 1.85 | 0.74 | 2.50 | 0.0123 |
| Obese vs. Normal | 4.69 | 0.70 | 6.74 | <0.0001 |
| **Time of Induction\*GDM:** |  |  |  |  |
| 4-8:GDM (Yes vs. No) | -10.00 | 3.47 | -2.88 | 0.0040 |
| 8-12:GDM (Yes vs. No) | -9.16 | 3.19 | -2.87 | 0.0042 |
| 12-16:GDM (Yes vs. No) | -9.07 | 2.36 | -3.84 | 0.0001 |
| 16-20:GDM (Yes vs. No) | -9.07 | 2.35 | -3.86 | 0.0001 |
| 20-24:GDM (Yes vs. No) | -7.54 | 2.57 | -2.93 | 0.0034 |

**Table 2.** Multiple comparisons of estimated marginal means of labor duration between GDM and non-GDM across different induction time intervals in the Sparrow cohort.

| Contrast (No vs. Yes) | Induction Time Interval | Estimate | SE | t.ratio | adj.p |
| --- | --- | --- | --- | --- | --- |
| No - Yes | 0-4 | -10.25 | 1.84 | -5.56 | <0.0001 |
| No - Yes | 4-8 | -0.25 | 2.94 | -0.09 | 0.9317 |
| No - Yes | 8-12 | -1.09 | 2.61 | -0.42 | 0.6760 |
| No - Yes | 12-16 | -1.18 | 1.48 | -0.79 | 0.4271 |
| No - Yes | 16-20 | -1.18 | 1.47 | -0.80 | 0.4227 |
| No - Yes | 20-24 | -2.71 | 1.79 | -1.51 | 0.1309 |

**Table 3.** Multiple comparisons of estimated marginal means of labor duration among different induction time intervals within either GDM or non-GDM in the Sparrow cohort.

| Contrast Among Induction Time Intervals | GDM | Estimate | SE | t.ratio | adj.p |
| --- | --- | --- | --- | --- | --- |
| (0-4) - (4-8) | No | 3.34 | 0.78 | 4.30 | 0.0001 |
| (0-4) - (8-12) | No | 1.61 | 0.66 | 2.44 | 0.0315 |
| (0-4) - (12-16) | No | 0.36 | 0.58 | 0.63 | 0.6623 |
| (0-4) - (16-20) | No | -0.18 | 0.58 | -0.31 | 0.8732 |
| (0-4) - (20-24) | No | -0.11 | 0.58 | -0.19 | 0.8945 |
| (4-8) - (8-12) | No | -1.73 | 0.79 | -2.19 | 0.0541 |
| (4-8) - (12-16) | No | -2.98 | 0.73 | -4.09 | 0.0002 |
| (4-8) - (16-20) | No | -3.52 | 0.73 | -4.82 | <0.0001 |
| (4-8) - (20-24) | No | -3.45 | 0.73 | -4.75 | <0.0001 |
| (8-12) - (12-16) | No | -1.24 | 0.60 | -2.07 | 0.0636 |
| (8-12) - (16-20) | No | -1.79 | 0.60 | -2.97 | 0.0091 |
| (8-12) - (20-24) | No | -1.72 | 0.60 | -2.87 | 0.0103 |
| (12-16) - (16-20) | No | -0.54 | 0.51 | -1.06 | 0.4350 |
| (12-16) - (20-24) | No | -0.47 | 0.51 | -0.93 | 0.4788 |
| (16-20) - (20-24) | No | 0.07 | 0.51 | 0.13 | 0.8945 |
| (0-4) - (4-8) | Yes | 13.33 | 3.38 | 3.95 | 0.0005 |
| (0-4) - (8-12) | Yes | 10.76 | 3.13 | 3.44 | 0.0022 |
| (0-4) - (12-16) | Yes | 9.43 | 2.29 | 4.12 | 0.0005 |
| (0-4) - (16-20) | Yes | 8.89 | 2.28 | 3.90 | 0.0005 |
| (0-4) - (20-24) | Yes | 7.43 | 2.50 | 2.97 | 0.0091 |
| (4-8) - (8-12) | Yes | -2.57 | 3.85 | -0.67 | 0.6046 |
| (4-8) - (12-16) | Yes | -3.90 | 3.21 | -1.22 | 0.4199 |
| (4-8) - (16-20) | Yes | -4.44 | 3.20 | -1.39 | 0.3538 |
| (4-8) - (20-24) | Yes | -5.91 | 3.36 | -1.76 | 0.1974 |
| (8-12) - (12-16) | Yes | -1.33 | 2.94 | -0.45 | 0.6985 |
| (8-12) - (16-20) | Yes | -1.87 | 2.93 | -0.64 | 0.6046 |
| (8-12) - (20-24) | Yes | -3.33 | 3.11 | -1.07 | 0.4725 |
| (12-16) - (16-20) | Yes | -0.54 | 2.02 | -0.27 | 0.7881 |
| (12-16) - (20-24) | Yes | -2.01 | 2.27 | -0.89 | 0.5639 |
| (16-20) - (20-24) | Yes | -1.47 | 2.26 | -0.65 | 0.6046 |

## DISCUSSION

Our study highlights the interplay between circadian rhythms and uterotonic responses to oxytocin in gravid mice, revealing that both the efficacy of oxytocin and the expression of oxytocin receptor (*Oxtr*) in the myometrium exhibit time-of-day-dependent variations. We report that drugs targeting the OXTR (the agonist oxytocin and the antagonist/inverse agonist atosiban) are markedly influenced by the time of day. These findings underscore the importance of considering time-of-day for studies on uterine contractility response to drugs. Moreover, our retrospective human data, show that time-of-the day of labor induction initiation play a major role in labor duration in women with and without diabetes. When labor induction was initiated around midnight in women with GDM, labor duration was significantly longer than in women without GDM. Independent of the presence or absence of GDM, labor duration was the shortest when labor induction was initiated in the early morning (4:00-8:00 hours). Together, these findings establish the circadian influence on drugs with uterotonic and tocolytic effects and make it imperative to consider the circadian system’s role in clinical studies as a potential strategy to reduce labor duration and improve maternal and neonatal outcomes.

### BMAL1 regulation of *Oxtr* expression defines the daily uterotonic efficacy of oxytocin in mice

The transcription factor BMAL1 is required for circadian rhythm generation throughout the body.^24,25,32^ In this study, we identify that *Oxtr* is under BMAL1-regulated transcriptional control *in vitro* and *in vivo*, which is associated with differential *Oxtr* expression in the myometrium. This finding contrasts with a previous study in this mouse model, which did not identify a difference in *Oxtr* expression in the *Bmal1* cKO uterus.^43^ This discrepancy could be due to the time of day of sample harvest, where we did identify a difference in *Oxtr-*expressing cells at ZT5. Importantly, we confirmed that the OXTR antagonist, atosiban, antagonized oxytocin-promoted uterine contractions at both studied time points; however, during the active phase of the day (ZT17), but not during the rest phase of the day (ZT5), atosiban significantly reduced baseline contraction frequency. This surprising finding may explain the variable efficacies in clinical studies using oxytocin antagonists to reduce uterine contractions and halt preterm labor.^48,49^ Given the recent withdrawal of the progestin Makena® from the US market and the current lack of FDA-approved tocolytics, establishing if atosiban might reliably halt preterm labor when given at specific times of the day, has the potential to have it reevaluated as a tocolytic in the US.

Despite extensive efforts, we could not validate a mouse OXTR antibody for western blotting or immunohistochemistry (not shown), impeding our capacity to demonstrate if the daily change in *Oxtr* mRNA translated to daily differences in protein expression and localization. Even though the protein studies failed, using *ex vivo* uterine contraction studies, we demonstrated that oxytocin more efficiently promoted uterine contractions during the active phase of the day (ZT17) in the GD18 control mouse. In addition, *Bmal1* cKO females had reduced uterotonic efficacy of oxytocin independent of the time-of-day. This reduction in oxytocin-promoted contractions in the *Bmal1* cKO matched the reduction, but not loss, of *Oxtr* in this mouse model. These results are consistent with studies highlighting BMAL1’s role in regulating circadian gene expression and its impact on physiological responses,^50^ and molecular clock-driven receptor expression and sensitivity to stimuli.^37^

### A mouse model of food-induced gestational diabetes (FID) shows an association of uterine molecular clock downregulation with reduced oxytocin-promoted contractions

While diabetes during pregnancy is known to deregulate the molecular clock in many tissues,^33,51,52^ and is associated with uterine dysfunctions, including diminished contractile response to Pitocin,^4^ our work is the first to demonstrate a correlation between myometrial molecular clock down-regulation and reduced uterine contractile response to oxytocin.

In mice, a high fat, high sucrose diet, independently of GDM, was associated with a significant reduction of *Clock* and *Bmal1* as compared to females on the standard chow diet (Ctr). Both FN (mice on high fat, high sugar diet which did not develop GDM) and FID groups presented with a reduction in the molecular clock. Still, only mice classified as having GDM, by failing the GTT (FID mice), became close to insensitive to the uterotonic effect of oxytocin at the studied time of day and the used oxytocin concentration (Figure 4E). This indicates a complex interaction between molecular clock expression and uterine sensitivity to oxytocin and highlights some significant differences between uterine function in mice on a high fat, high sucrose diet with and without GDM. A recent study reported similar data demonstrating that mice on a long-term high-caloric diet causing obesity significantly reduced uterine contractions before and during labor *in vivo*.^53^ It is important to note that our study, which was shorter in duration for the time on the high-calorie diet than the other study, did not lead to the development of obesity (Supplemental Figure 5A). In addition, there were differences in experimental designs between the two studies hindering direct comparisons. Of note, the previous study identified an association between uterine contractile malfunction in food-induced obesity mice and increased medium and long-chain fatty acid uptake in the myometrium.^53^ This finding is consistent with previous research indicating that metabolic disorders can disrupt circadian rhythms.^54,55^ However, unlike our study, diet-induced obesity did not significantly reduce *Oxtr*, *Bmal1*, *Clock*, *Cry2,* or *Oxtr* expression in the GD18 uterus (not shown, GSE268397),^53^ a difference potentially linked to different diets and durations of diets used in the two studies. Both mouse studies demonstrated the established poor contractile function in diabetic women; however, our model also recapitulated the reduced uterotonic efficacy in women with GDM.^4^ Together, these results show a differential impact of diet and diabetes on uterine contractile function, and further studies are needed to understand the molecular mechanisms involved in diet and GDM-induced changes in uterine contractile function.

### The time-of-day of induction of labor initiation defines labor duration in a GDM-dependent manner

GDM is associated with a higher risk for pregnancy complications than non-diabetics, including prolonged labor, preterm birth, abnormal uterine contractions, emergency cesarean section, and postpartum hemorrhage.^3–6^ While there is evidence that spontaneous uterine contractions and sensitivity to oxytocin *in vivo* changes depending on the time of day,^56,57^ little is known about what regulates this daily change in uterine function. While there is evidence that spontaneous uterine contractions and sensitivity to oxytocin *in vivo* changes depending on the time of day,^13,56–58^ little is known about what regulates this daily change in uterine function. In agreement with this prior work, we here show that mouse and human myometrial cells have defined daily time windows of increased and decreased contractile response to oxytocin. Our previous work in a larger human cohort with nulli-, primi-, and multiparous women without diabetes identified a circadian rhythm to labor duration based on the time of day of initiating labor induction.^14^ Here, we extend this line of inquiry by focusing on women with GDM, where labor duration is significantly longer than non-diabetics in our two independent datasets. Focusing on the Sparrow cohort, which was considerably larger than the McLaren cohort, we found that during induction of labor when labor induction was initiated around midnight (0:00-4:00 hours), labor duration was significantly longer in women with diabetes (GDM yes), than non-diabetic women (GDM no). Importantly, we found that the shortest labor duration was obtained independently of the presence of GDM when labor induction was initiated at 4:00-8:00 hours (early morning). Although the McLaren dataset did not reach statistical significance when split into 4-hour bins, this dataset trended in the same direction as the Sparrow dataset. Together, these data identify a potential benefit in considering the time of day of labor induction for women with and without GDM, and specifically show that inducing women with diabetes close to midnight (0:00-4:00 hours) should be avoided due to the significantly longer labor duration resulting from induction at this time of day.

### Conclusion

Our findings identify that the time-of-day of oxytocin in mice, or labor induction in women defines uterine contraction frequency in mice and labor duration in women, respectively. Using the mouse as a model, we show that *Bmal1* in the myometrium drives daily changes in *Oxtr* expression and oxytocin efficacy. Importantly, we show that food-induced gestational diabetes (FID) dampens the uterotonic efficacy of oxytocin in the mouse at the studied time point. These mouse studies translated to women, where the time-of-day of labor induction impacted labor duration in women with and without GDM. Specifically, GDM was associated with significantly longer labor durations when labor induction was initiated around midnight. Altogether, this work adds to a growing literature showing how the time of day defines drug efficacy,^59,60^ and highlights the importance of time-of-day for initiation of labor induction in normal pregnancies, and pregnancies complicated by gestational diabetes.

## MATERIALS AND METHODS

### Mouse breeding and timed matings

Sex as a biological variable was not considered in this study. Pregnant female mice were used for all experiments. Male mice were exclusively used for mating purposes. Mice were maintained on a light/dark cycle of 12 h light, 12 h dark, with lights ON (150-300 lux in the cage) at Zeitgeber Time 0 (ZT0) and lights OFF at ZT12. Mice had food and water ad libitum. All mice were from a C57BL/6 genetic background 5-14 weeks-of-age at the start of experiments and 12-24 weeks-of-age at time of euthanasia. *PER2::Luciferase* [Tg(Per2-luc)1Jt, JAX #006852] and *Bmal1^flox^* [B6.129S4(Cg)-Arntl^tm1Weit^/J, JAX #007668] were crossed with *Telokin^Cre^*^+/-^ mice,^43^ the *Bmal1^flox/flox^:Telokin^Cre+/-^* mice are referred to as *Bmal1* cKO. We confirmed the specific deletion of *Bmal1* in smooth muscle (Supplemental Figure 6). Mice with germline recombination were excluded from the study. For timed matings, the day after vaginal plug identification was considered as GD1.

### Food-induced gestational diabetes mouse model

Controls were maintained on standard chow for the entire study. To generate mice with food-induced gestational diabetes (FID), 5-6-week-old females had blood glucose levels monitored using One Touch Verio Blood Glucose Monitoring System Meter (Meijer) and body weight measured pre- and post-conception. After basal glucose levels were determined, the mice were given a high (45%) fat diet (TEKLAD: TD.06415), with 30% corn syrup in autoclaved water. Mice were maintained on this diet for 5 weeks (Figure 4A). After 5 weeks, both control and high fat, high sugar diet females were mated with a male on standard chow. The day of the presence of a vaginal plug was defined at GD1. A glucose tolerance test (GTT) was done on GD16. Briefly, pregnant females were starved from ZT0-ZT6. A bolus of sucrose (2g/kg weight) was given through intraperitoneal injection and blood glucose was monitored at 0’, 15’, 30’, 60’, 90’, and 120’. Mice were euthanized by cervical dislocation on GD19 at ZT5±2 hours, and uterine samples were prepared for uterine contraction studies. Pups were sexed, weighed and had insulin and blood glucose measured at euthanasia. The high fat, high sucrose diet-induced gestational diabetes in 60% of the mice (FID), whereas 40% of the females did not develop diabetes (FN).

### Organotypic circadian PER2::Luciferase luminescence recording

PER2::luciferase (PER2::LUC) recordings were done as we have previously described.^32^ In brief, the uterus was removed from a GD18, ZT19±3h mouse, and uterine samples were collected. For samples with endometrium removed, the endometrium was gently scraped from the myometrium with a scalpel. Uterine explants were placed on a MilliCell membrane in 35-mm culture dishes and placed into a LumiCycle (Actimetrics) at 35.5°C, 5%/95% CO_2_, in a non-humidified environmental chamber. The bioluminescence signal was counted every 10 minutes for 6 days and analyzed on days 1–6 of recording time. Data were normalized by subtraction of the 24-hour running average from the raw data and then smoothed with a 1 hour running average (Luminometer Analysis, Actimetrics) and analyzed blind to the experimental group. PER2::LUC recordings were analyzed by the Luminometer Analysis software (Actimetrics) with LM fit (damped sin) as the mathematical model.

### Mouse uterine contractions

GD18 females were euthanized by cervical dislocation at ZT5±3h or ZT17±3h. The uterus was dissected along the longitudinal axis into full-thickness uterine strips (endometrium + myometrium) of ∼ 2×5 mm^2^ in ice cold oxygenated physiological saline solution [154 mM NaCl, 5.6 mM KCl, 1.2 mM MgSO_4_, 10.2 mM HEPES, 2 mM CaCl_2_ and 8 mM glucose]. The uterine strips were mounted in tissue organ baths (DMT Muscle Strip Myograph System-820MS) containing 6 mL oxygenated physiological saline solution at 36.0 ± 0.5°C to mimic physiological conditions. Each strip was stretched to a final tension of 6 mN. Uterine strips were allowed to equilibrate for ∼2 hours until 15 minutes of regular spontaneous contractions occurred. The tissues were then treated with vehicle (Milli Q water, at 1/2000 dilution), 500 nM atosiban (A3480- 10MG, Sigma Life Science) or 1 nM oxytocin (O3251-1000IU, Sigma Life Science). At the end of all recordings, uterine strips were washed 3x with 6 mL physiological saline solution, 10 minutes between washes, and then treated with 100 mM KCl. Treatment with 100 mM KCl was used to confirm uterine viability. LabChart software (ADInstruments) analyzed contractions by evaluating area AUC, amplitude and frequency of contractions over a period of 10 minutes during baseline or drug application periods.

### Quantitative Real-Time PCR

For RT-qPCR, the full-thickness uterine strips were first dissected from pregnant females at designated time points on GD19 as described above and immediately frozen and stored at −80°C until RNA extraction. Total RNA was extracted using E.Z.N.A. ® Total RNA Kit I (Omega Bio-tek). cDNA was obtained by reverse transcription of total RNA using an Lunascript® RT Supermix cDNA synthesis kit (New England Biolabs). cDNA products were detected using an iQ SYBR Green Supermix (Bio-Rad Laboratories) on a qRT-PCR CFX real-time detection system (Bio-Rad Laboratories). qRT-PCR primers were on *Bmal1* (F: TGACCCTCATGGAAGGTTAGAA; R: GGACATTGCATTGCATGTTGG), *Clock* (F: ATGGTGTTTACCGTAAGCTGTAG; R: CTCGCGTTACCAGGAAGCAT), and *Cry2* (F: CACTGGTTCCGCAAAGGACTA; R: CCACGGGTCGAGGATGTAGA). Data were expressed as fold change using the ΔΔCT method by normalizing *Bmal1, Clock, and Cry2* to housekeeping *H2afz (*F: TCACCGCAGAGGTACTTGAG; R: GATGTGTGGGATGACACCA) and *Gapdh* (F: GGCAAATTCCATGGCACCGT; R: GCAAATGAGCCCCAGCCTTC). Data represent mean fold change ± SEM.

### Transient transfections and luciferase assays

Mouse NIH3T3 cell culture, transient transfections and site-directed mutagenesis was completed as we have previously described^61,62^ and are described in Supplemental Figure 7.

### *In vitro* PHM1-41 cell contraction study

Immortalized pregnant human myometrium cells from 39 weeks gestation, termed PHM1-41^47^ (American Type Culture Collection, VA, #CRL-3046), were cultured in DMEM (Mediatech) containing 10% fetal bovine serum (Gemini Bio), and 1x penicillin-streptomycin (Life Technologies/Invitrogen) in a humidified 5% CO_2_ incubator at 37°C. PHM1-41 cells were seeded into 35cm plates (Nunc) at 18 million cells per well. To synchronize the molecular clock, cells were first treated with low serum media [DMEM containing 0.1% bovine serum albumin (BP9706-100, Fisher)] for 24 hours (serum starvation, Figure 5A).^45,46^ After 24 hours serum starvation, cells were then treated with regular cell culture media. Beginning at 12 hours after synchronization (set as circadian time point 00:00 hours), cells were treated at the indicated time-points with either vehicle (water) or 4.5 nM oxytocin (O3251-1000IU, Sigma Life Science). Images of cells were captured at 0 and 15 minutes after treatment with a Leica DMi1 microscope (Leica) under bright field at 20x magnification. Images were analyzed for cellular surface area as a measure of contraction using Fiji.^63^ Percent cell size was calculated as cellular surface area at 15 minutes divided by the corresponding cellular surface area at baseline (0 minutes). For each image 5-10 cells were analyzed.

### RNAscope multiplex *in situ* hybridization assay

We completed RNAscope® multiplex *in situ* hybridization detection of mouse (*Mus musculus*) mRNAs with RNAscope® LS Multiplex Fluorescent Reagent Kit (Advanced Cell Diagnostics, cat no. 322800) following vendor’s standard protocol for FFPE tissue sections with minor modifications. RNAscope® assays were performed on a Leica Bond autostainer as described^64,65^ with the following probes: RNAscope® 2.5 LS Probe – Mm-Arntl (also known as Bmal1) [aryl hydrocarbon receptor nuclear translocator-like (Arntl) transcript variant 1 mRNA, cat no. 438748- C1] and RNAscope® 2.5 LS Probe – Mm- Oxtr-C2 [oxytocin receptor (Oxtr), cat no. 412178-C2]. Tissue slides were counterstained with DAPI and scanned with Aperio Versa imaging system with 20X objective with customized narrow-width band excitation and emission filter cubes as described.^64^ The Aperio Cellular IF Algorithm (Leica Biosystems, No: 23CIFWL) was used for automated cell enumeration and segmentation based on nuclear DAPI staining. Cells were classified based on the expression levels of one or more mRNAs (cell classes were threshold based solely on increasing intensity value per pixel within each segmented object/cell).

### Human participants

Sex as a biological variable was not considered. Only pregnant females were enrolled in this study. Two independent cohorts, the Sparrow cohort (n= 7,804 term deliveries at the Sparrow Hospital, East Lansing, MI, from February 2019 to March 2022) and the McLaren cohort (n= 1,632 term deliveries at the McLaren Greater Lansing, MI, from December 2019 to December 2023), were used. Deidentified data on pregnant women was collected from the electronic medical records at the two hospitals. All patients with maternal age <18 years old, gestational age <37 weeks, multiple birth=Yes, and induction=No or NA were excluded. This exclusion led to two final datasets with 3613 and 587 patients from the Sparrow and McLaren cohorts, respectively, for downstream analyses.

#### Measurements of major variables

GDM patients were defined as those with ICD10 code = O24 in Sparrow or comorbidity codes = ‘diabetes’ or ‘gestational diabetes’ in McLaren, and the remaining patients as Non-GDM (including those with missing values (NAs)) in both cohorts. Induction of labor duration was calculated by subtracting time-of-induction (defined by the initial dose of oxytocin or any cervical ripening agent, such as foley catheter or cytotec/misoprostol) from delivery time. Labor induction start time was grouped into six intervals – 0:00-4:00, 4:00- 8:00, 8:00-12:00, 12:00-16:00, 16:00-20:00, and 20:00-24:00 hours. Body Mass Index (BMI) was grouped into 3 groups – Normal (BMI= 18.5 to 25 kg/m^2^), Overweight (BMI= 25 to 30 kg/m^2^), and Obese (higher than 30 kg/m^2^). There were no underweight subjects. Maternal age was categorized as <30 versus ≥30 years old. Parity in the Sparrow cohort was dichotomized into “0” versus “≥1”. Gravidity in McLaren (parity information was not available) was grouped into “1” versus “>1”. Race in Sparrow was coded as: “White”, “Hispanic”, “Black”, “Asian”, and “Others”. Ethnicity in McLaren was coded as “ Hispanic” and “ Non-Hispanic” due to the information about race being unavailable.

#### Statistical analysis of retrospective data

All categorical variables were summarized as numbers and percentages, and continuous variables (i.e., labor duration) were present as mean and standard deviation (SD). Bivariate associations between labor duration and individual covariates were analyzed using one-way ANOVA (for at least 3 groups) or two-sample t-tests (for two groups). A multiple linear regression model was applied to examine the association between outcome–labor duration and multiple covariates with an interaction term of induction time*GDM. Comparisons of estimated marginal means among interactions of factors after adjusting the means of other factors in the multiple linear regression model were conducted with the emmeans R package.^66^ The adjusted p-values were calculated by using the Benjamini-Hochberg method to correct multiple comparisons. All data management and analyses were conducted in R environment (version 4.4.1).^67^

### Study approval

All animal procedures were performed according to the protocols approved by the Animal Use Committee and the Institutional Animal Care of Michigan State University and conducted by the Guide for the Care and Use of Laboratory Animals (National Research Council, 2011). The Institutional Review Board of Michigan State University approved human research under STUDY0007199 (Sparrow Health System) and STUDY00010164 (McLaren Greater Lansing).

### Statistical analysis

*Ex vivo* uterine contractions of both control and *Bmal1* cKO mice were analyzed with One or Two-way ANOVA. Other statistical analyses were done as described in the figure legends. All data management and statistical analyses were done using Prism 9 (GraphPad Software, San Diego, CA, USA), R (R Development Core Team).

## Data availability

Data, mouse models, code, and human datasets will be made available upon request and approval by the institutions.

## Author contributions

The experimental design was done by T.V.D., A.M.Y., G.Z., N.S., A.S.C., A.S., L.S., I.N.O, R.L., R.S. and H.M.H.; data collection was by T.V.D., A.M.Y., N.S., A.S.C., D.N, N.L.; data analysis was completed by T.V.D., A.M.Y., G.Z., N.S., A.S.C., N.L, A.S., L.S., I.N.O, R.L., R.S. R.C.and H.M.H.; and all the authors did data interpretation and manuscript writing and approval.

## Acknowledgments

This work was funded by the USDA National Institute of Food and Agriculture Hatch project MICL1018024 (H.M.H.), March of Dimes Grant no 5-FY19-111 (H.M.H.), the NIH/National Institute of Environmental Health Sciences project R01ES035691 (H.M.H.), and a sub-award from the Michigan Diabetes Research Center through the National Institute of Diabetes and Digestive and Kidney Diseases P30DK020572 (H.M.H.). A.M.Y. was supported by the Eunice Kennedy Shriver National Institute of Child Health & Human Development (NICHD) of the National Institutes of Health under Award Numbers F32HD107852 and K99HD113843 with additional support of training by T32HD087166. We thank Angela Smet, MD, and Rachel Eck, for feedback and help with data analysis; Dr. Louis Muglia at Cincinnati Children’s Hospital Medical Center in Cincinnati (Ohio, USA) for sharing the Telokin^Cre^ mice. We also thank MSU Precision Health Program Tissue Analysis Core for technical assistance. We acknowledge the Obstetrics Initiative (OBI) for providing access to the McLaren data used in this study. Support for OBI is provided by Blue Cross Blue Shield of Michigan and Blue Care Network as part of the BCBSM Value Partnerships program.

## Notes

### Competing Interest Statement

The authors have declared no competing interest.

