## Supplemental methods and data for "Interaction between time-of-day and oxytocin efficacy in mice and humans with and without gestational diabetes"

### Cell culture conditions, transient transfections and luciferase assays

Mouse embryonic fibroblast NIH-3T3 (American Type Culture Collection, VA, #CRL-1658) cells were cultured in DMEM (Mediatech), containing 10% fetal bovine serum (Gemini Bio), and 1x penicillin-streptomycin (Life Technologies/Invitrogen) in a humidified 5% $CO_2$ incubator at 37˚C. For luciferase assays, NIH3T3 cells were seeded into 24-well plates (Nunc) at 50,000 cells per well. Transient transfections for luciferase assays were performed using PolyJet™ (SignaGen Laboratories), following the manufacturer's recommendations. NIH3T3 cells were co-transfected as indicated in the figure legends with 200 ng/well luciferase reporter plasmids, as well as 100 ng/well thymidine kinase-β-galactosidase reporter plasmid, which served as an internal control.[1,2] The plasmids used were mouse -1000 to +200bp Oxtr-luciferase (200 ng/uL, VectorBuilder), mouse Bmal1 overexpression plasmid (200 ng/uL (Addgene, #31367), pcDNA 3.1 (between 0ng/uL and 200 ng/uL), and pGL4-luciferase (200 ng/uL). Site-directed mutagenesis of the Oxtr-luciferase reporter was performed using the NEB Q5 Site-Directed Mutagenesis Protocol (New England Biolabs Inc.), following manufacturer's instructions, to mutate each E'-box to TGACGA. Primers for NEB Q5 site-directed mutagenesis were designed using NEBaseChanger[3] (Supplemental Figure 7). We systematically equalized plasmid concentrations by adding the corresponding inactive plasmid backbone to equalize the amount of DNA transfected into cells. Cells were given their respective ligands 24 h after transfection and were then harvested 48 h after transfection in lysis buffer [100 mM potassium phosphate (pH 7.8) and 0.2% Triton X-100]. Luciferase values were normalized to β-galactosidase values to control transfection efficiency. Values were further normalized by expression as fold change compared to pGL4 control plasmid, as indicated in the figure legends.

**Supplemental figures**

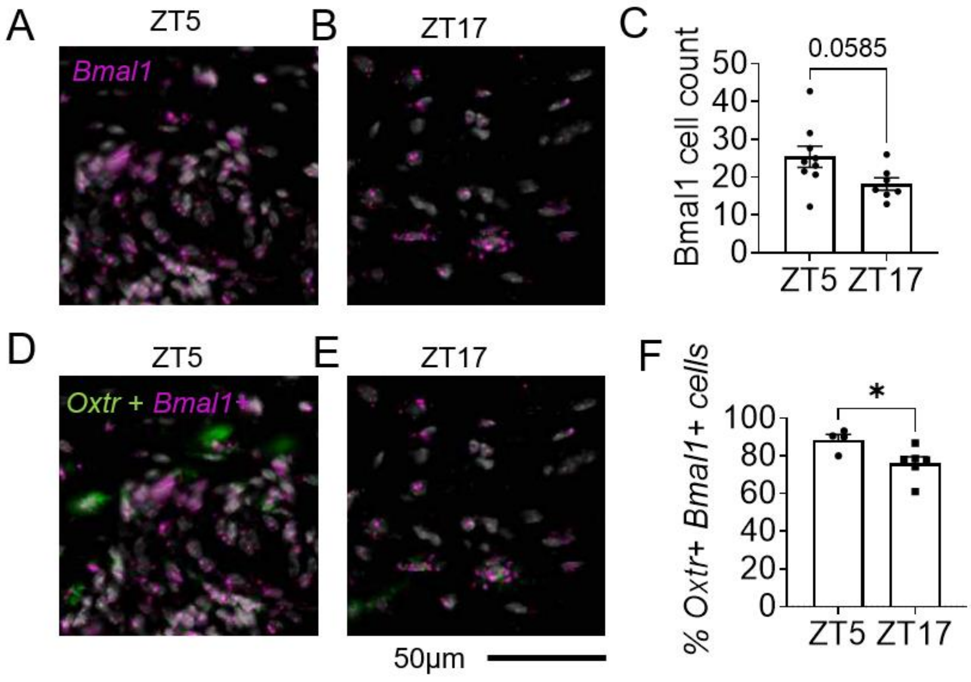


**Supplemental Figure 1.** Time of day defines myometrial *Bmal1 and Oxtr* expression. Image of ISH RNAscope of GD18 uterus at **A**) ZT5 and **B**) ZT17 with Green: *Oxtr*; violet: *Bmal1;* white: DAPI. **C**) Histogram of cell counts expressing *Bmal1.* T-test, n = 7-9/group. Image of ISH RNAscope of GD18 uterus at **D**) ZT5 and **E**) ZT17 and **F**) histogram of the percentage of cells expressing *Oxtr* and *Bmal1.* T-test, n = 4-5/group. *, p<0.05.

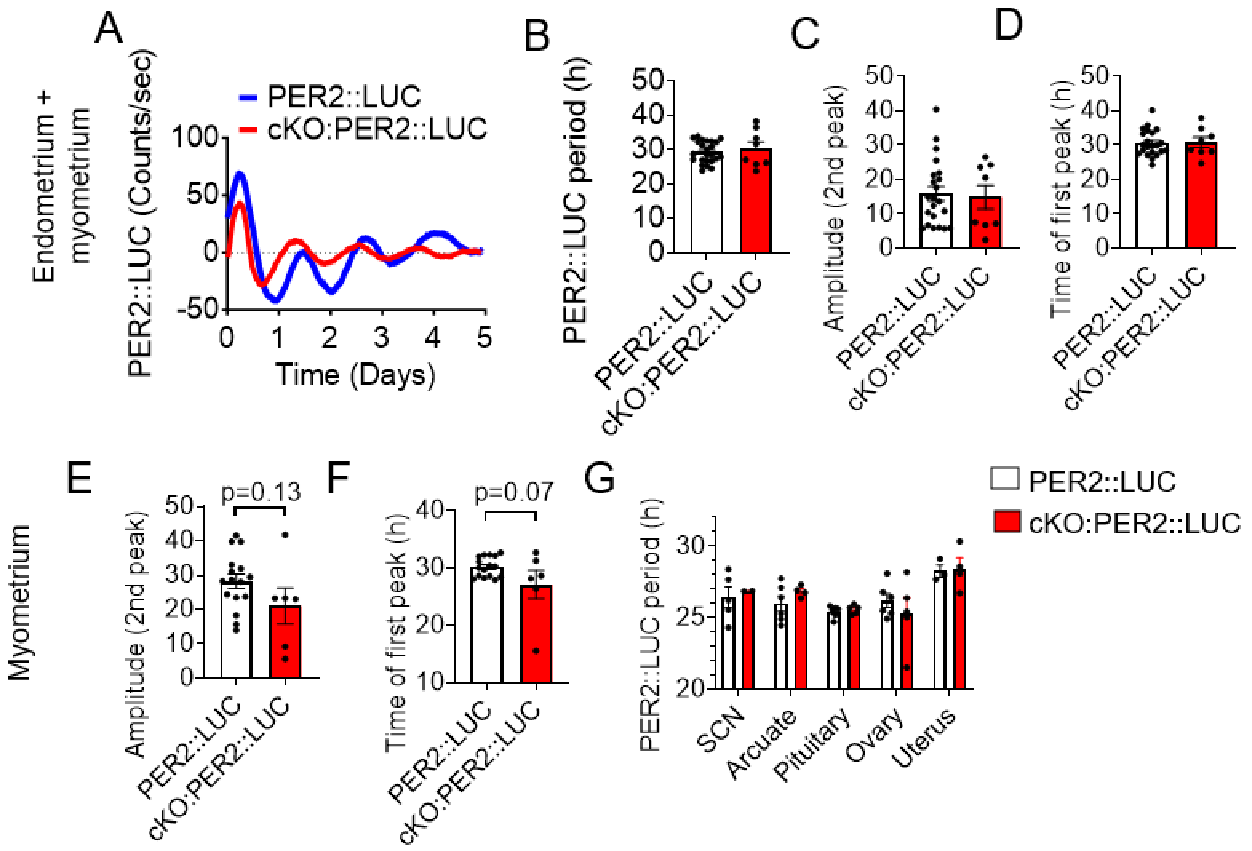


**Supplemental Figure 2.** The myometrium from GD18 female mice exhibit circadian rhythms that are abolished without BMAL1. **A**) Full-thickness GD18, ZT19 uterine explants from PER2::LUCIFERASE reporter mice (PER2::LUC, blue) and triple transgenic cKO:PER2::LUC mice (red) displayed cyclic PER2::LUCIFERASE (PER2::LUC) recordings over 6 days. Comparing **A-D**) full-thickness uterine explants and **E-F**) endometrium removed (myometrium) uterine explants, between control (PER2::LUC, blue) and *Bmal1* cKO (cKO:PER2::LUC, red). **B**) PER2::LUC period (hours) **C, E**) amplitude of the second peak of PER2::LUC rhythms and **D, F**) time of day of the first peak of PER2::LUC rhythms. **G**) To assess the impact of *Bmal1* deletion in the myometrium on circadian timekeeping in the reproductive axis, the period of PER2::LUC expression was evaluated in tissues harvested at ZT19 suprachiasmatic nucleus (SCN), arcuate nucleus (arcuate), pituitary, ovary and full-thickness uterine explants (uterus) in GD18 control (PER2::LUC) and *Bmal1* cKO (cKO:PER2::LUC). Statistical analysis (B-F) Student's t-test, n=2-7 per group, and (G) two-way ANOVA, n=2-6.

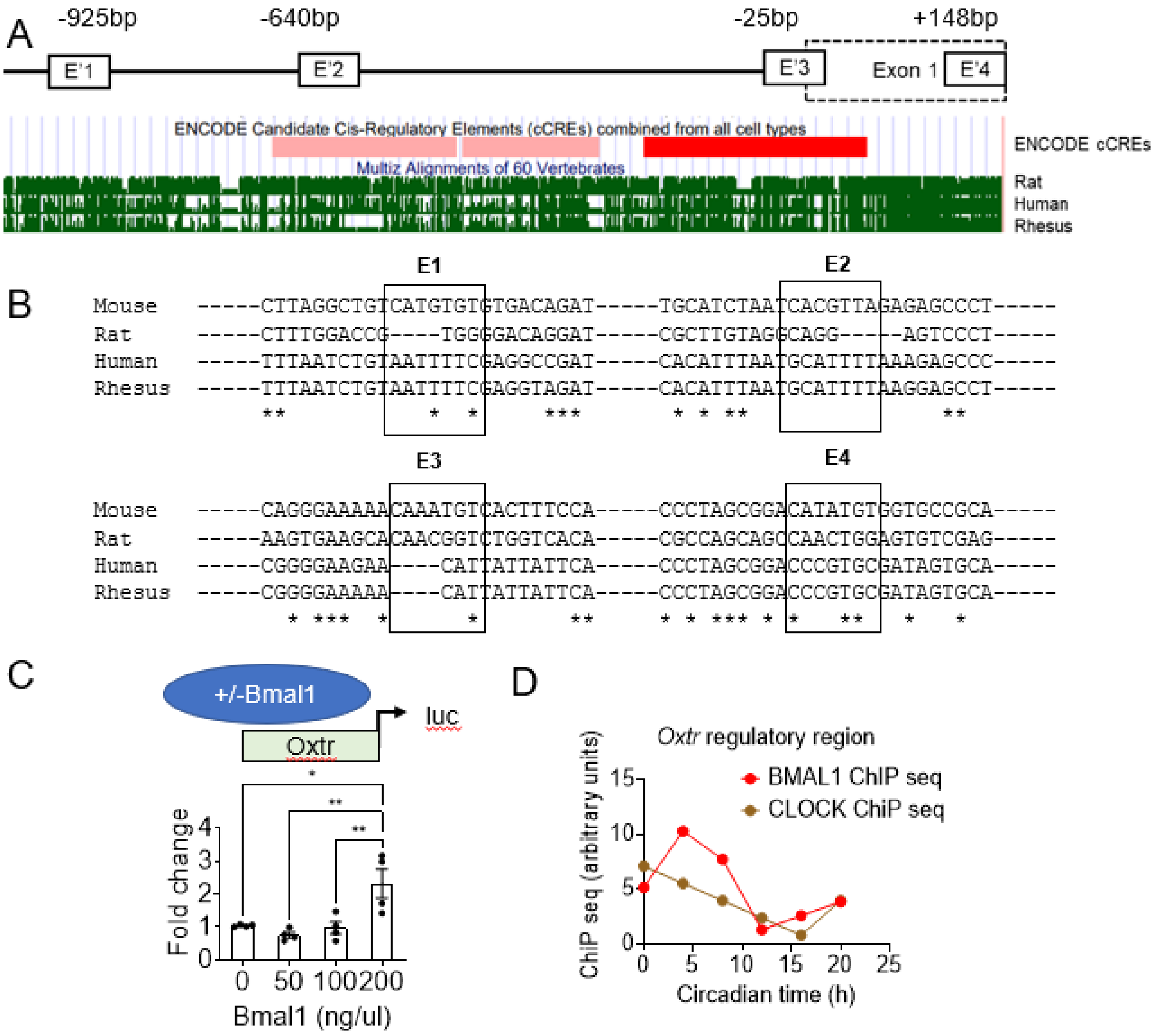


**Supplemental Figure 3**. **A**) DNA sequence alignment was done with Clustal Omega and the UCSC Genome Browser. Alignment with Clustal Omega was done on sequences from -1000bp to +200bp from the beginning of the *Oxtr* genes of mouse, rat, human and *Rhesus* monkey obtained from NCBI (August 2021). **B**) Clustal Omega identified four semi-conserved BMAL1 binding sites (E'-boxes) in the *Oxtr* regulatory region (E1, E2, E3, E4). The * indicates conserved nucleotides. **C**) To determine if BMAL1 regulated *Oxtr* expression *in vitro*, NIH-3T3 cells were transiently transfected with mouse *Bmal1* overexpressing plasmid and mouse *Oxtr*-luciferase plasmids. One-way ANOVA, n=4 per group, in duplicate **D)** CHiP seq of BMAL1 and CLOCK binding to the regulatory region of the mouse *Oxtr* in the adult male liver. Data extracted from Koike *et al.* Science 2012. [4]

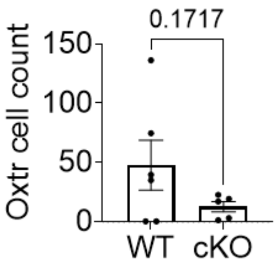


**Supplemental Figure 4.** Quantified ISH RNAscope for *Oxtr* expressing cells in the GD18 control (WT) and *Bmal1* cKO (cKO) uterus at ZT19 +/- 3h. n = 5-6/group. T-test.

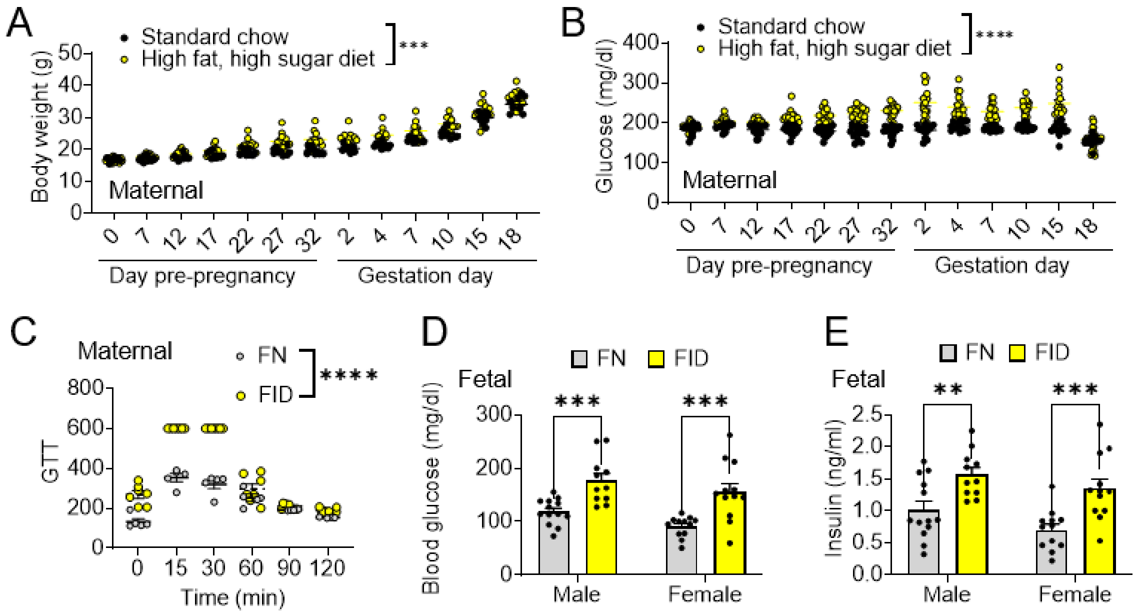


**Supplemental Figure 5. Maternal and fetal characteristics from the high-fat high sugar diet study**. Maternal **A**) weight and **B)** blood glucose during the experimental period for Control (Ctr) mice on standard chow and mice on high fat, high sucrose diet, n= 16/group. **C**) Glucose tolerance test (GTT) on gestation day 16 (GD16). Females were separated into high-fat, high sugar diet with-normal GTT (FN) and high-fat, high sugar diet-induced gestational diabetes (FID) based on peak glucose levels in response to the GTT. n= 5-7. GD19 male and female blood **D)** glucose level and **E)** insulin levels. N=11-13/group. Two-way ANOVA, ****, p<0.0001.

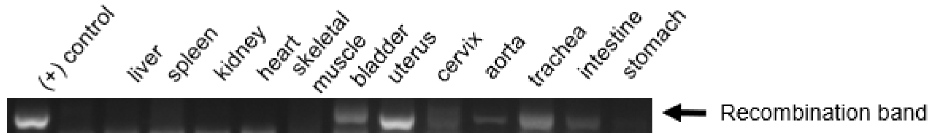


**Supplemental Figure 6.** PCR to identify Telokin-cre recombination of the Bmal-flox/flox allele in the mouse. Indicated tissues were collected from adult female *Bmal1* cKO and analyzed for *Bmal1*$^{flox/flox}$-allele recombination. The presence of a white band indicates Cre-induced recombination.

| E'-box | Original sequence | Actual mutated sequence | Forward primer<br>(Uppercase = target-specific primer) | Reverse primer |
|---|---|---|---|---|
| E1 | CATGTG | TGACGA | CTTAGGCTGTtgacgaTGTGACAGATACTAATG | GTTACAAAACACTCAGGTC |
| E2 | CACGTT | GACG-A | TGCATCTAATtgacgaAGAGAGCCCTG | TGTTTTATCCTGACATGTTATTTC |
| E3 | CATATG | TCGTCA | CCCTAGCGGAtgacgaTGGTGCCGCAGCTCAGGGTTC | GGCAGAGCAAACCGGCCG |
| E4 | CAAATG | TCGTCA | CAGGGAAAAAtgacgaTCACTTTCCAAGGTTCCTATATCTCTG | AGTAAATTGTAATAAAGACGC |

**Supplemental Figure 7.** E'-box sequences and primers for site-directed mutagenesis.

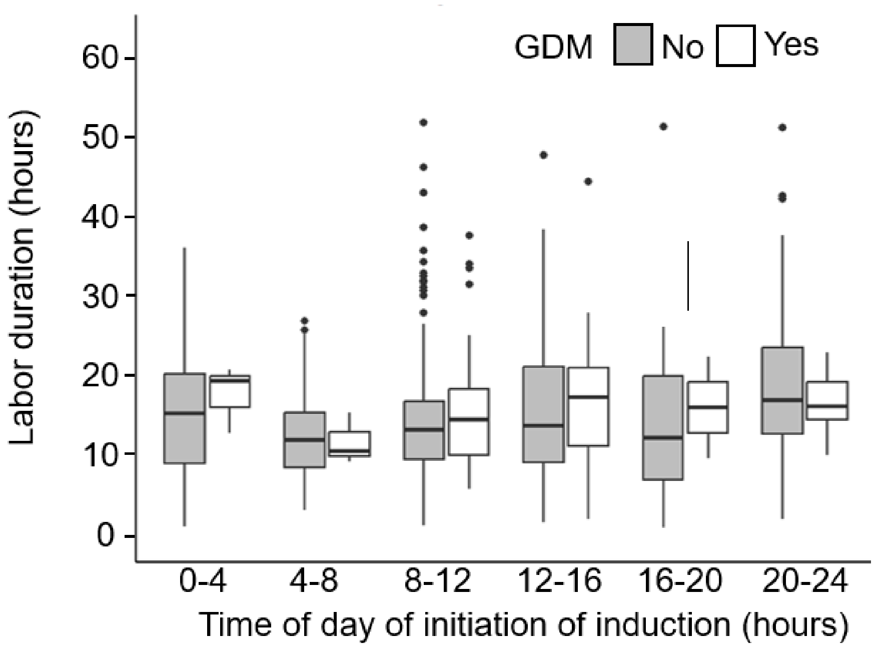


**Supplemental Figure 8. Labor duration as a function of the time of day of labor induction in women with and without GDM.** The average labor duration for each 4 hour time of induction bin across the 24 hour day is presented for women with (Yes, n=56) and without (No, n=531) GDM in the McLaren cohort. Linear regression with a comparison of labor duration as a function of time of induction, with and without GDM. $p>0.05$. For complete statistics, see Tables S4 and S5.

**Table S1.** Frequency distribution of the patients across all covariates (Sparrow, N=3613)

| | | Duration of Labor | | |
|---|---|---|---|---|
| | n (%) | Mean (SD) | t or F Statistic | p |
| **Race:** | | | | |
| White | 2457 (68.0) | 18.3 (11.1) | 0.45 | 0.7740 |
| Hispanic | 189 (5.2) | 18.3 (10.6) | | |
| Black | 528 (14.6) | 18.6 (11.7) | | |
| Asian | 132 (3.7) | 17.2 (10.5) | | |
| Others | 307 (8.5) | 18.2 (10.5) | | |
| **Maternal Age:** | | | | |
| <30 | 1990 (55.1) | 19.1 (11.4) | 4.72 | <0.0001 |
| ≥30 | 1623 (44.9) | 17.4 (10.7) | | |
| **Parity:** | | | | |
| 0 | 1727 (47.8) | 23.0 (11.7) | 26.74 | <0.0001 |
| ≥1 | 1886 (52.2) | 14.0 (8.4) | | |
| **BMI:** | | | | |
| Normal (BMI<25) | 223 (6.2) | 14.3 (8.3) | 41.60 | <0.0001 |
| Overweight (BMI 25-30) | 980 (27.1) | 16.5 (10.3) | | |
| Obese (BMI>30) | 2410 (66.7) | 19.5 (11.4) | | |
| **Time of Induction:** | | | | |
| 0-4 | 515 (14.3) | 19.3 (12.0) | 5.81 | <0.0001 |
| 4-8 | 259 (7.2) | 15.8 (9.0) | | |
| 8-12 | 446 (12.3) | 17.0 (9.7) | | |
| 12-16 | 807 (22.3) | 18.1 (10.9) | | |
| 16-20 | 790 (21.9) | 18.7 (11.5) | | |
| 20-24 | 796 (22.0) | 19.1 (11.5) | | |
| **GDM:** | | | | |
| Yes | 187 (5.2) | 20.6 (13.0) | 2.92 | 0.0035 |
| No | 3426 (94.8) | 18.2 (11.0) | | |

**Table S2.** Frequency distribution of the patients across all covariates (McLaren, N=587)

| | n (%) | Duration of Labor Mean (SD) | t or F Statistic | p |
|---|---|---|---|---|
| **Ethnicity:** | | | | |
| Hispanic | 25 (5.1) | 14.3 (8.2) | -0.25 | 0.8003 |
| Non-Hispanic | 463 (94.9) | 14.7 (7.8) | | |
| **Maternal Age:** | | | | |
| <30 | 407 (69.3) | 14.4 (7.7) | -1.61 | 0.1087 |
| ≥30 | 180 (30.7) | 15.6 (8.4) | | |
| **Gravidity:** | | | | |
| 1 | 419 (71.4) | 14.6 (7.6) | -0.70 | 0.4872 |
| >1 | 168 (28.6) | 15.1 (8.7) | | |
| **BMI:** | | | | |
| Normal (BMI<25) | 162 (32.3) | 13.3 (7.1) | 5.50 | 0.0043 |
| Overweight (BMI 25-30) | 133 (26.5) | 15.1 (8.3) | | |
| Obese (BMI>30) | 206 (41.1) | 16.0 (8.2) | | |
| **Time of Induction:** | | | | |
| 0-4 | 34 (5.8) | 15.3 (8.2) | 5.25 | 0.0001 |
| 4-8 | 32 (5.5) | 13.4 (5.2) | | |
| 8-12 | 268 (45.7) | 13.5 (7.0) | | |
| 12-16 | 117 (19.9) | 15.2 (8.5) | | |
| 16-20 | 55 (9.4) | 15.2 (9.1) | | |
| 20-24 | 81 (13.8) | 18.4 (8.9) | | |
| **GDM:** | | | | |
| Yes | 56 (9.5) | 15.8 (7.5) | 1.02 | 0.3100 |
| No | 531 (90.5) | 14.7 (8.0) | | |

**Table S3**. Summary of the association between the outcome variable Labor Duration and the covariates (McLaren, N=501).

| | Coefficient (β) | SE of β | t Statistic | p |
|---|---|---|---|---|
| **Intercept:** | 14.47 | 1.57 | 9.21 | <0.0001 |
| **Time of Induction:** | | | | |
| 4-8 vs. 0-4 | -1.98 | 2.14 | -0.93 | 0.3540 |
| 8-12 vs. 0-4 | -2.66 | 1.59 | -1.67 | 0.0955 |
| 12-16 vs. 0-4 | -0.75 | 1.72 | -0.43 | 0.6636 |
| 16-20 vs. 0-4 | -0.73 | 1.89 | -0.39 | 0.6989 |
| 20-24 vs. 0-4 | 2.45 | 1.79 | 1.37 | 0.1709 |
| **GDM:** | | | | |
| Yes vs. No | 0.05 | 1.19 | 0.04 | 0.9682 |
| **BMI*:** | | | | |
| Overweight vs. Normal | 1.95 | 0.92 | 2.12 | 0.0341 |
| Obese vs. Normal | 2.65 | 0.83 | 3.18 | 0.0016 |
| **Time of Induction*GDM:** | | | | |
| 4-8:GDM (Yes vs. No) | -5.26 | 9.77 | -0.54 | 0.5908 |
| 8-12:GDM (Yes vs. No) | -2.72 | 5.92 | -0.46 | 0.6465 |
| 12-16:GDM (Yes vs. No) | -2.87 | 6.26 | -0.46 | 0.6475 |
| 16-20:GDM (Yes vs. No) | 0.80 | 6.99 | 0.11 | 0.9085 |
| 20-24:GDM (Yes vs. No) | -5.79 | 6.76 | -0.86 | 0.3924 |

* n=86 missing values.

**Table S4**. Multiple comparisons of estimated marginal means of labor duration between GDM and non-GDM across different induction time intervals in the McLaren cohort.

| Contrast (No vs. Yes) | Induction Time Interval | Estimate | SE | df | t.ratio | adj.p |
|---|---|---|---|---|---|---|
| No - Yes | 0-4 | -2.76 | 5.71 | 487 | -0.48 | 0.6290 |
| No - Yes | 4-8 | 2.50 | 7.94 | 487 | 0.32 | 0.7529 |
| No - Yes | 8-12 | -0.04 | 1.60 | 487 | -0.03 | 0.9792 |
| No - Yes | 12-16 | 0.11 | 2.61 | 487 | 0.04 | 0.9673 |
| No - Yes | 16-20 | -3.56 | 4.06 | 487 | -0.88 | 0.3808 |
| No - Yes | 20-24 | 3.03 | 3.63 | 487 | 0.83 | 0.4048 |

**Table S5**. Multiple comparisons of estimated marginal means of labor duration among different induction time intervals within either GDM or non-GDM in the McLaren cohort.

| Contrast Among Induction Time Intervals | GDM | Estimate | SE | df | t.ratio | adj.p |
|---|---|---|---|---|---|---|
| (0-4) - (4-8) | No | 1.98 | 2.14 | 487 | 0.93 | 0.5900 |
| (0-4) - (8-12) | No | 2.66 | 1.59 | 487 | 1.67 | 0.2389 |
| (0-4) - (12-16) | No | 0.75 | 1.72 | 487 | 0.44 | 0.7488 |
| (0-4) - (16-20) | No | 0.73 | 1.89 | 487 | 0.39 | 0.7488 |
| (0-4) - (20-24) | No | -2.45 | 1.79 | 487 | -1.37 | 0.3204 |
| (4-8) - (8-12) | No | 0.68 | 1.62 | 487 | 0.42 | 0.7488 |
| (4-8) - (12-16) | No | -1.24 | 1.74 | 487 | -0.71 | 0.6981 |
| (4-8) - (16-20) | No | -1.25 | 1.91 | 487 | -0.66 | 0.6981 |
| (4-8) - (20-24) | No | -4.43 | 1.81 | 487 | -2.45 | 0.0731 |
| (8-12) - (12-16) | No | -1.91 | 1.01 | 487 | -1.90 | 0.1724 |
| (8-12) - (16-20) | No | -1.93 | 1.27 | 487 | -1.52 | 0.2774 |
| (8-12) - (20-24) | No | -5.11 | 1.11 | 487 | -4.59 | 0.0001 |
| (12-16) - (16-20) | No | -0.02 | 1.43 | 487 | -0.01 | 0.9896 |
| (12-16) - (20-24) | No | -3.20 | 1.29 | 487 | -2.48 | 0.0731 |
| (16-20) - (20-24) | No | -3.18 | 1.50 | 487 | -2.11 | 0.1319 |
| (0-4) - (4-8) | Yes | 7.24 | 9.53 | 487 | 0.76 | 0.8211 |
| (0-4) - (8-12) | Yes | 5.38 | 5.70 | 487 | 0.94 | 0.8211 |
| (0-4) - (12-16) | Yes | 3.61 | 6.02 | 487 | 0.60 | 0.8211 |
| (0-4) - (16-20) | Yes | -0.07 | 6.74 | 487 | -0.01 | 0.9912 |
| (0-4) - (20-24) | Yes | 3.34 | 6.52 | 487 | 0.51 | 0.8211 |
| (4-8) - (8-12) | Yes | -1.86 | 7.92 | 487 | -0.24 | 0.9395 |
| (4-8) - (12-16) | Yes | -3.63 | 8.16 | 487 | -0.44 | 0.8211 |
| (4-8) - (16-20) | Yes | -7.32 | 8.70 | 487 | -0.84 | 0.8211 |
| (4-8) - (20-24) | Yes | -3.90 | 8.52 | 487 | -0.46 | 0.8211 |
| (8-12) - (12-16) | Yes | -1.77 | 2.88 | 487 | -0.61 | 0.8211 |
| (8-12) - (16-20) | Yes | -5.45 | 4.17 | 487 | -1.31 | 0.8211 |
| (8-12) - (20-24) | Yes | -2.04 | 3.80 | 487 | -0.54 | 0.8211 |
| (12-16) - (16-20) | Yes | -3.69 | 4.60 | 487 | -0.80 | 0.8211 |
| (12-16) - (20-24) | Yes | -0.27 | 4.28 | 487 | -0.06 | 0.9912 |
| (16-20) - (20-24) | Yes | 3.42 | 5.22 | 487 | 0.65 | 0.8211 |

**Supplemental data references**


1. Hoffmann HM, Trang C, Gong P, Kimura I, Pandolfi EC, Mellon PL. Deletion of Vax1 from gonadotropin-releasing hormone (GnRH) neurons abolishes GnRH expression and leads to hypogonadism and infertility. *J Neurosci*. 2016. doi:10.1523/JNEUROSCI.2723-15.2016

2. Meadows JD, Breuer JA, Lavalle SN, et al. Deletion of Six3 in post-proliferative neurons produces weakened SCN circadian output, improved metabolic function, and dwarfism in male mice. *Mol Metab*. 2022;57:101431. doi:10.1016/j.molmet.2021.101431

3. Hoffmann HM, Gong P, Tamrazian A, Mellon PL. Transcriptional interaction between cFOS and the homeodomain-binding transcription factor VAX1 on the GnRH promoter controls Gnrh1 expression levels in a GnRH neuron maturation specific manner. *Mol Cell Endocrinol*. 2018;461:143-154. doi:10.1016/j.mce.2017.09.004

4. Koike N, Yoo SH, Huang HC, et al. Transcriptional architecture and chromatin landscape of the core circadian clock in mammals. *Science (80- )*. 2012;338(6105). doi:10.1126/science.1226339
